# Interleukin-11 is a Marker for Both Cancer- and Inflammation-Associated Fibroblasts that Contribute to Colorectal Cancer Progression

**DOI:** 10.1101/2020.01.25.919795

**Authors:** Takashi Nishina, Yutaka Deguchi, Wakami Takeda, Masato Ohtsuka, Daisuke Ohshima, Soh Yamazaki, Mika Kawauchi, Eri Nakamura, Chiharu Nishiyama, Yuko Kojima, Satomi Adachi-Akahane, Mizuho Hasegawa, Mizuho Nakayama, Masanobu Oshima, Hideo Yagita, Kazutoshi Shibuya, Tetuo Mikami, Naohiro Inohara, Norihiro Tada, Hiroyasu Nakano

## Abstract

Interleukin (IL)-11 is a member of the IL-6 family of cytokines and involved in multiple cellular responses, including tumor development. However, the origin and functions of IL-11-producing (IL-11^+^) cells are not fully understood. To characterize IL-11^+^ cells *in vivo*, we generated *Il11* reporter mice. IL-11^+^ cells appeared in the colon of three murine tumor models, and a murine acute colitis model. *Il11ra1* or *Il11* deletion attenuated the development of colitis-associated colorectal cancer. IL-11^+^ cells expressed fibroblast markers, and genes associated with cell proliferation and tissue repair. IL-11 induced STAT3 phosphorylation in colonic fibroblasts, suggesting the activation of IL-11^+^ fibroblasts. Analysis using the human cancer database revealed that genes enriched in IL-11^+^ fibroblasts were elevated in human colorectal cancer, and correlated with reduced disease-free survival. Together, our results suggested that tumor cells induced IL-11^+^ fibroblasts, and that a feed-forward loop between IL-11 and IL-11^+^ fibroblasts might contribute to tumor development.

## INTRODUCTION

Maintenance of intestinal homeostasis involves a variety of cell types, including epithelial, immune, and stromal cells^1,2,3^. Within the intestinal lamina propria, stromal cells include fibroblasts, α smooth muscle actin (αSMA)-positive myofibroblasts, endothelial cells, and pericytes^4, 5^. These stromal cells organize the tissue architecture, and have recently been revealed to play crucial roles in regulating immune responses, tissue repair, and tumor development^3,4,5^. Recent studies have focused on fibroblasts that can support tumor growth, termed cancer-associated fibroblasts (CAFs)^6,7,8^. In a very recent study, single-cell RNA-sequencing (scRNA-seq) was performed to analyze colon biopsies from healthy individuals and ulcerative colitis (UC) patients. The results revealed that UC patients’ colon samples included a unique subset of fibroblasts, termed inflammation-associated fibroblasts (IAFs), with high expression of *IL11*, *IL24*, *IL13RA2,* and *TNFSFR11B*^9^. Another prior study demonstrated that CD10 and GPR77 might be markers of CAFs that mediate chemotherapy resistance in human cancer through IL-6 and IL-8 production^10^. Combined these findings together, stromal fibroblasts play a crucial role in the development of cancer and colitis, the full picture of fibroblast function and heterogeneity remains unclear.

Interleukin (IL)-11 is a member of the IL-6 family, and exhibits pleiotropic functions, including hematopoiesis, bone development, tissue repair, and tumor development^11, 12^. The IL-11 receptor comprises IL-11Rα1, which binds IL-11, and gp130, which transmits signals to the nucleus via Janus kinase (JAK) activation^13^. JAKs phosphorylate STAT3, and phosphorylated STAT3 enters the nucleus where it activates the transcription of various target genes associated with cell proliferation and apoptosis suppression^14,15,16,17^. IL-11 production is regulated by several cytokines, including TGFβ, IL-1β, IL-17A, and IL-22^18, 19, 20, 21^. We previously demonstrated that IL-11 production is induced by reactive oxygen species (ROS) and the electrophile 1,2-naphthoquinone, which, in turn, promotes liver and intestine tissue repair^22, 23^. While IL-11 is reportedly produced in various cell types (including stromal, hematopoietic, and epithelial cells) in response to different stimuli^16, 17, 24, 25^, the cellular sources of IL-11 *in vivo* are not fully understood.

Colorectal cancers exhibit increased IL-11 expression in human and mice. Moreover, deletion of the *Il11ra1* gene attenuates colitis-associated colorectal cancer (CAC) development in mice treated with azoxymethane (AOM) and dextran sulfate sodium (DSS), or in adenomatous polyps of mice harboring mutation of the *Adenomatous polyposis coli (Apc)* gene (*Apc^Min/+^*)^17^. On the other hand, *IL11* gene polymorphism is associated with increased ulcerative colitis (UC) susceptibility in human patients ^26^. *IL11* expression is increased in patients with mild UC, but decreased in patients with severe UC^27^. These findings suggest that IL-11 may promote colorectal cancer development, but could also attenuate colitis under certain conditions. Moreover, recent studies report that IL-11 plays a crucial role in fibrosis development in various organs, including the lung, heart, liver, and kidney^28, 29^.

In the present study, we aimed to characterize IL-11-producing cells *in vivo*. We generated *Il11*-*Enhanced green fluorescence protein (Egfp)* reporter mice, and detected IL-11^+^ cells in the colonic tumor tissues of a CAC mouse model and in *Apc^Min/+^* mice. Deletion of *Il11ra* or *Il11* attenuated CAC development in mice. When tumor organoids were transplanted into the colon, IL-11^+^ cells appeared in the tumor tissues. Moreover, IL-11^+^ cells rapidly appeared in the colon of mice after DSS treatment. IL-11^+^ cells expressed stroma cell markers and genes associated with cell proliferation and tissue repair, suggestive of colonic fibroblasts. IL-11 induced robust STAT3 phosphorylation in colonic fibroblasts, but not in colonic epithelial organoids. Using the human cancer database, we found that the genes enriched in IL-11^+^ fibroblasts were elevated in human colorectal cancer, and that high expression of several of these genes was correlated with reduced disease-free survival among CRC patients. Together, our present results demonstrated that tumor cells induced IL-11^+^ fibroblasts, and that a feed-forward loop between IL-11 and IL-11^+^ fibroblasts may contribute to tumor development.

## RESULTS

### Characterization of IL-11^+^ Cells in CAC Using *Il11-Egfp* Reporter Mice

To characterize IL-11-producing (IL-11^+^) cells *in vivo*, we generated transgenic mice in which *Egfp* expression was under control of the *Il11* gene promoter using a bacterial artificial chromosome (BAC) vector (Supplementary Fig. 1a). *Egfp* cDNA and a polyA signal were inserted in-frame in the second exon of the *Il11* gene (Supplementary Fig. 1a). As expected, *Il11* mRNA expression was correlated with *Egfp* mRNA expression in various tissues (Supplementary Fig. 1b, c). Notably, *Il11* mRNA expression was highest in the testis, and was very low in other tissues in mice under normal conditions (Supplementary Fig. 1b).

Although *Il11* mRNA expression is elevated in colorectal cancer among mice and human^16, 17^, the origin of IL-11^+^ cells remains controversial. We used the *Il11-Egfp* reporter mice to monitor the appearance of IL-11^+^ cells in a CAC model. AOM administration followed by repeated exposure to DSS causes CAC development in mice^30,31,32,33^. On day 77 after AOM/DSS administration, the *Il11-Egfp* reporter mice developed large tumors in the colon (Fig. 1a, b). At this time, we isolated tumor and nontumor tissues, and examined the *Il11* mRNA expression by qPCR. *Il11* and *Egfp* mRNA expressions were elevated in tumor tissues compared to nontumor tissues from mice with AOM/DSS-induced CAC (Fig. 1c).

**Fig. 1.**
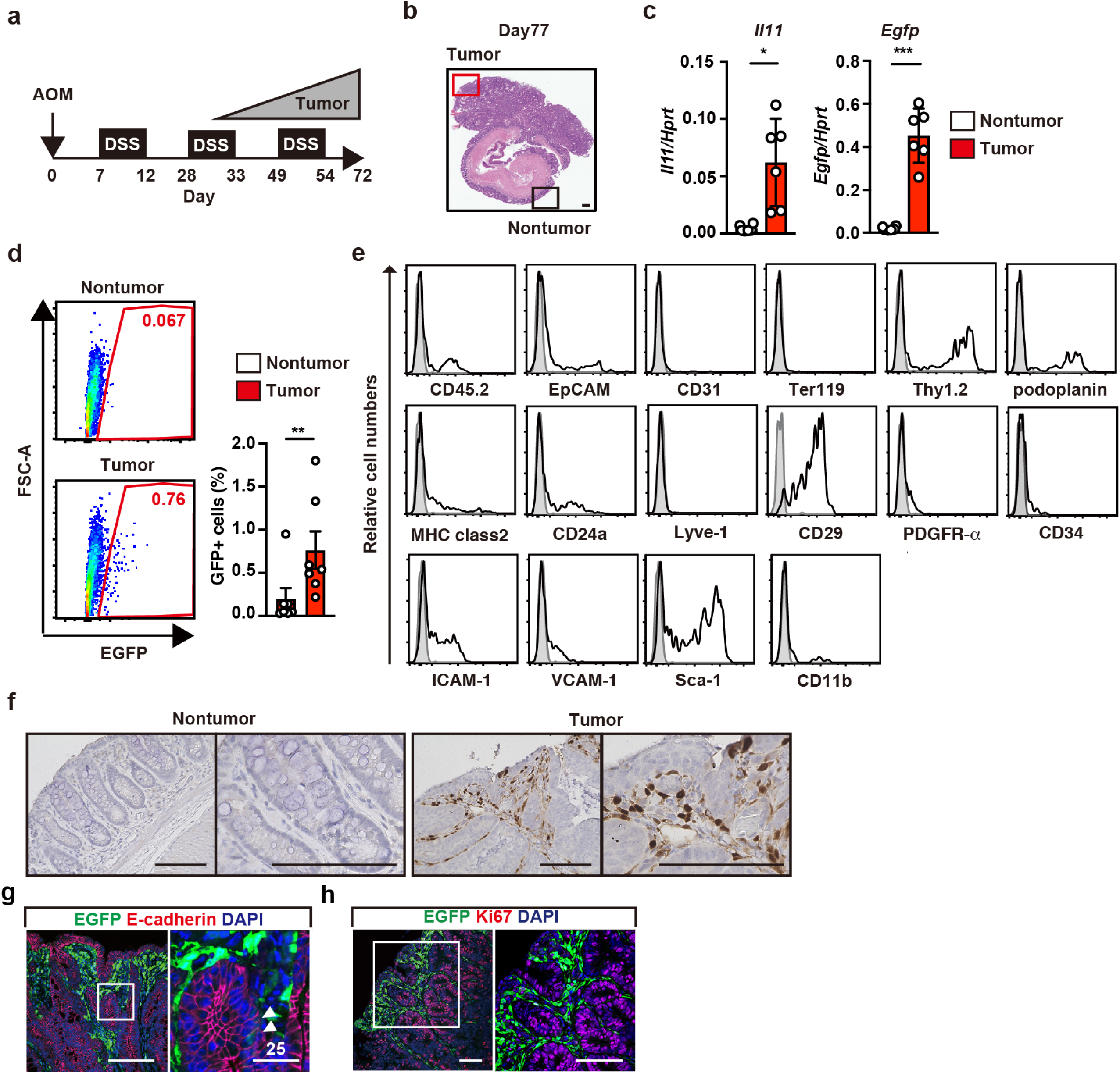
Characterization of IL-11^+^ Cells in CAC Using *Il11-Egfp* Reporter Mice. **a** Protocol for induction of AOM/DSS-induced CAC in mice. *Il11-Egfp* reporter mice were intraperitoneally injected with AOM on day 0, followed by repeated DSS administration. Colorectal cancer gradually develops on ∼30 days after AOM injection. Unless otherwise indicated, the following experiments used tumor and nontumor tissues collected on day 98–105 after AOM/DSS treatment. **b** Representative image of tumor and adjacent nontumor tissues in the colon of mice on day 77 after AOM injection. Colonic sections were stained with hematoxylin & eosin (H&E). Scale bar, 200 μm. **c** Prepared mRNA samples from tumor and nontumor tissues were analyzed by qPCR to determine expression of the indicated genes. Results are mean ± SE (n = 6–10 mice). **d, e** From colonic tumor tissues of wild-type or *Il11-Egfp* reporter mice, single-cell suspensions were prepared, and percentages of IL-11^+^ cells were determined. Representative flow cytometry images are shown (**d**). Results are mean ± SE (n = 7 mice). Cells were stained with the indicated antibodies, and expression on EGFP^+^ cells was analyzed by flow cytometry (**e**). Results are representative of four independent experiments. **f** Nontumor and tumor tissue sections were stained with anti-GFP antibody. Scale bars, 100 μm. **g, h** Tumor sections were stained with the indicated antibodies (red) and with anti-GFP antibody (green) (n = 3 mice). Right panels show enlarged images of white boxes from left panels. White arrowheads indicate merged cells. Scale bars, 100 μm, unless otherwise indicated. Statistical significance was determined by two-tailed unpaired Student’s *t*-test (**c, d**). *p < 0.05; **p < 0.01; ***p < 0.001.

We next isolated and characterized IL-11^+^ cells from tumors of *Il11-Egfp* reporter mice. We observed increased numbers of EGFP (IL-11)^+^ cells in tumor tissues compared to nontumor tissues from the colon of AOM/DSS-treated *Il11-Egfp* reporter mice (Fig. 1d). While the majority of IL-11^+^ cells expressed stroma cell markers, such as Thy1.2 and podoplanin, numerous IL-11^+^ cells expressed CD45.2 or EpCAM (Fig. 1e). Immunohistochemistry (IHC) revealed IL-11^+^ cells in stroma tissues surrounding tumor cells (Fig. 1f). IL-11^+^ cells also expressed vimentin, collagen I, and collagen IV, but not αSMA (Supplementary Fig. 1d), suggesting that these cells were fibroblasts but not myofibroblasts. Notably, a few E-cadherin-positive tumor cells exhibited weak EGFP expression (Fig. 1g). Overall, these results suggest that the IL-11^+^ cells constituted heterogenous cell populations, which might explain previous apparently inconsistent results showing that IL-11^+^ cells were derived from hematopoietic, epithelial, or stromal cells^17, 24^.

EGFP-positive cells were not detected in the colon of *Il11-Egfp* reporter mice before AOM/DSS treatment (data not shown), possibly suggesting that small numbers of resident IL-11^+^ cells might proliferate and expand *in situ*. Alternatively, IL-11^−^ cells could be converted into IL-11^+^ cells after AOM/DSS treatment. To discriminate these two possibilities, we investigated whether IL-11^+^ cells expressed the cell-proliferating antigen Ki67. The majority of IL-11^+^ cells did not express Ki67 (Fig. 1h), supporting the possibility that IL-11^+^ cells were derived from IL-11^−^ cells during tumor development.

### Attenuated CAC Development in *Il11ra1^−^*^/*−*^ and *Il11^−^*^/*−*^ Mice

A previous study reported attenuated CAC development in *Il11ra1 ^−^* ^/*−*^ mice^17^. Consistently, our present data confirmed the attenuation of AOM/DSS-induced CAC in *Il11ra1^−^*^/*−*^ mice compared to wild-type mice (Supplementary Fig. 2a). To further substantiate that the IL-11/IL-11R-dependent signaling pathway contributed to CAC development, we investigated *Il11^−^*^/*−*^ mice. As we previously reported^34^, *Il11^−^*^/*−*^ mice showed no abnormalities. When wild-type and *Il11^−^*^/*−*^ mice were treated with AOM/DSS, we observed attenuated CAC development in *Il11^−^*^/*−*^ mice compared to wild-type mice (Supplementary Fig. 2b). Moreover, reciprocal BM transfer experiments revealed that *Il11* expression in non-hematopoietic cells might be primarily responsible for the increased tumor numbers and tumor load in the colon (Supplementary Fig. 2c, d), although *Il11* expression in hematopoietic cells also partly contributed the increased tumor area in the colon (Supplementary Fig. 2c, d). We focused our subsequent analyses on IL-11^+^ colonic fibroblasts, which are hereafter referred to as IL-11^+^ colon cancer-associated fibroblasts (CAFs).

### IL-11^+^ CAFs Appear in Tumor Tissues in the Absence of Inflammation

The above-described results suggested that tumor cells may have educated the surrounding IL-11*^−^* cells to become IL-11^+^ CAFs. An alternative possibility is that IL-11*^−^* cells might cell-autonomously become IL-11^+^ CAFs within the setting of an inflammatory milieu triggered by DSS administration. To discriminate these possibilities, we used two different inflammation-independent murine tumor models: adenomatous polyps in *Apc^Min/+^* mice^33^ and transplantation of tumor organoids into wild-type mice. In *Apc^Min/+^* mice, *Il11* and *Egfp* mRNA expression levels were elevated in colon tumors compared to nontumor tissues (Fig. 2a), and IL-11^+^ CAFs appeared in stroma tissues surrounding tumor cells in the colon (Fig. 2b). Consistent with the results of IHC, we found increased numbers of IL-11^+^ CAFs in the tumor tissues compared to non-tumor tissues from the colon of *Apc^Min/+^* mice (Fig. 2c). Moreover, most IL-11^+^ CAFs expressed podoplanin and Thy1.2, and small numbers of IL-11^+^ cells expressed hematopoietic and epithelial cell markers (Fig. 2d). Again, we observed weak EGFP expression in some E-cadherin-positive tumor cells themselves (Fig. 2e). Notably, most IL-11^+^ CAFs were not Ki67^+^ (Fig. 2f). IL-11^+^ CAFs expressed vimentin, collagen I, and collagen IV, but not αSMA (Fig. 2g).

**Fig. 2.**
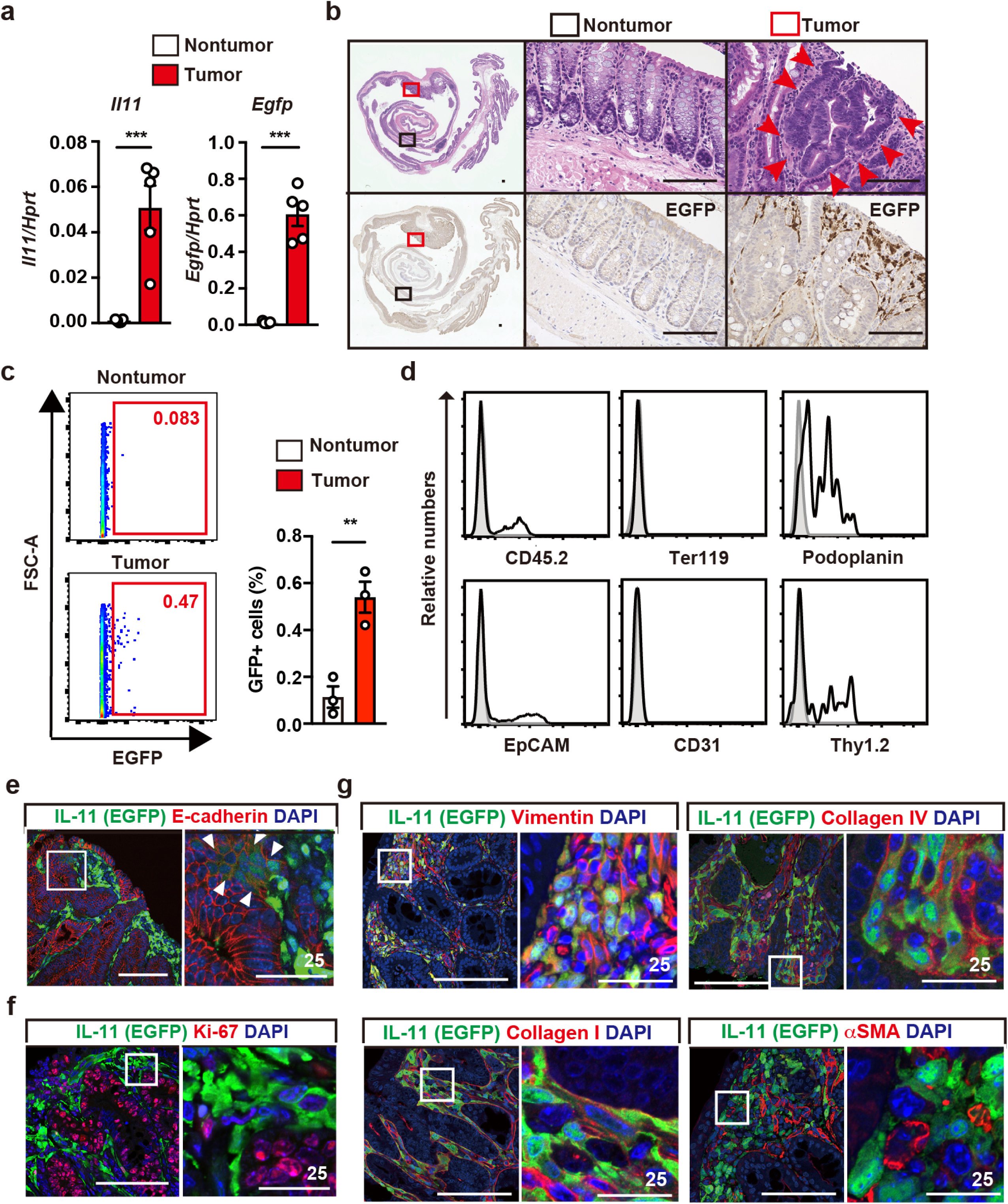
IL-11^+^ Cells Appear in Tumor Tissues in the Absence of Inflammation. **a** Elevated *Il11* and *Egfp* expression in colonic tumors from *Apc^min/+^;Il11-Egfp* reporter mice. Experiments were performed using tumor and nontumor tissues from the colon of 20- to 24-week-old *Apc^min/+^;Il11-Egfp* reporter mice. *Il11* and *Egfp* mRNA expression was determined by qPCR. Results are mean ± SE (n = 5 mice). **b** Colonic tissue sections from *Apc*^min/+^;*Il11-Egfp* reporter mice were stained with H&E (upper panels) or anti-GFP antibody (lower panels). Middle and right panels, respectively, show enlargements of the black and red boxes from the left panels. Red arrowheads indicate tumor cells. Scale bar, 100 μm. **c, d** IL-11^+^ cells express stromal cell markers and, to a lesser extent, hematopoietic or epithelial cell markers. Colonic cells were prepared from tumors of *Apc*^min/+^;*Il11-Egfp* reporter mice, stained with the indicated antibodies, and analyzed by flow cytometry. Percentages of IL-11^+^ cells were determined (**c**). Results are mean ± SE (n = 3 mice). Representative histograms show expressions of the indicated markers on EGFP^+^ cells (**d**). Results are representative of three independent experiments. **e–g** Characterization of IL-11^+^ cells by IHC. Colonic tumor sections from *Apc*^min/+^;*Il11-Egfp* reporter mice were stained with the indicated antibodies (red) and anti-GFP antibody (green). White arrowheads indicate merged cells. Scale bar, 100 μm, unless otherwise indicated. Statistical significance was determined by two-tailed unpaired Student’s *t*-test (**a, c**). **p < 0.01; ***p < 0.001.

We previously generated tumor organoids from the intestines of AKTP mice, which exhibit mutations of *Apc, Kras, Tgfbr2,* and *Tp53* in intestinal epithelial cells ^35^. These tumor organoids were transferred into the colon of *Il11-Egfp* reporter mice, and we examined whether IL-11^+^ CAFs appeared along with tumor development (Supplementary Fig. 3a). On day 30 post-transplantation, the tumor organoids had developed into large tumors in the colon and, importantly, IL-11^+^ CAFs appeared in the tumor tissues (Figure Supplementary Fig. 3b). Moreover, these IL-11^+^ CAFs expressed podoplanin and vimentin, but not CD45 or E-cadherin (Supplementary Fig. 3c). Together, these results strongly support that tumor cells instructed IL-11^−^ cells to become IL-11^+^ CAFs in the absence of inflammation.

### IL-11^+^ Cells Appear in the Colon of DSS-Treated Mice and Express Stromal Cell Marker

To test whether inflammation alone induced IL-11^+^ cell development, we treated *Il11-Egfp* reporter mice with only DSS. *Il11* and *Egfp* expressions were very low in the colon of untreated *Il11-Egfp* reporter mice, and were elevated in the colon of *Il11-Egfp* reporter mice on day 7 after DSS treatment (Fig. 3a). IHC revealed that on day 6 after DSS treatment, large numbers of IL-11^+^ cells appeared in the subepithelial tissues, where intestinal epithelial cells were detached due to severe inflammation (Fig. 3b). Moreover, we detected the rapid appearance of IL-11^+^ cells just one day after DSS treatment (Fig. 3c). For a more detailed comparison of the phenotypes of IL-11^+^ CAFs in tumor tissues and the IL-11^+^ cells that appeared in colitis, we used flow cytometry to analyze the expressions of various cell surface markers. After DSS treatment, we observed increased numbers of IL-11^+^ cells in the colon (Fig. 3d). These IL-11^+^ cells expressed stroma cell markers, including Thy1.2, podoplanin, CD29, PDGFR-α, ICAM-1, VCAM-1, and Sca-1, but not CD45.2, EpCAM, CD31, or Ter119 (Fig. 3e). IL-11^+^ cells were also positive for vimentin, collagen I, and collagen IV, but not αSMA (Fig. 3f), suggesting that these cells were fibroblasts, but not myofibroblasts. Moreover, these IL-11^+^ cells did not incorporate 5-bromo-2ʹ-deoxyuridine (BrdU), a hallmark of cell proliferation, suggesting that they did not proliferate *in situ* (Fig. 3g). Together, these findings suggest that inflammation alone was sufficient to induce IL-11^+^ cell development.

**Fig. 3.**
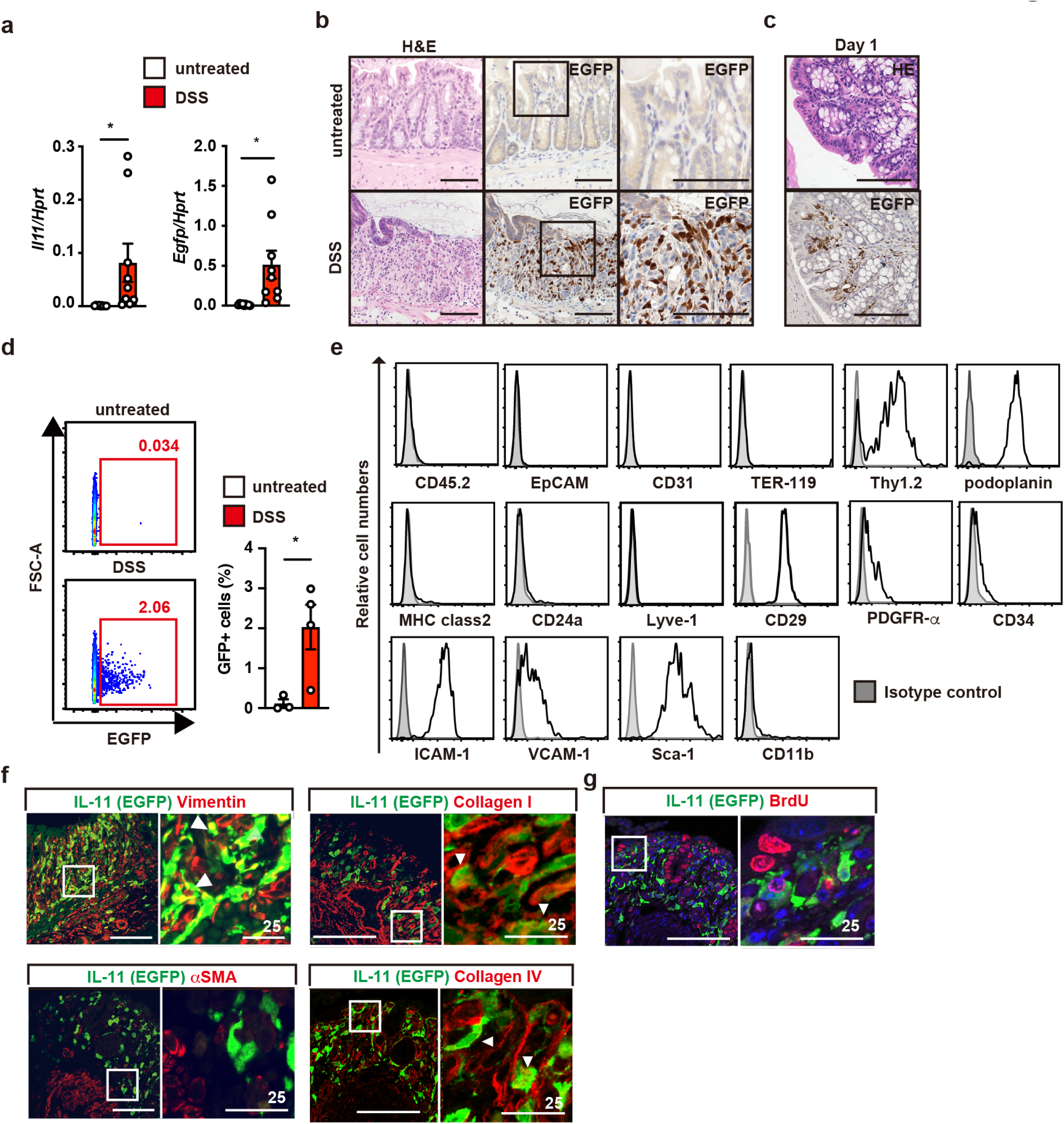
IL-11^+^ Cells Appear in the Colon of DSS-Treated Mice and Express Stromal Cell Marker. **a** *Il11-Egfp* reporter mice were treated with 1.5% DSS in drinking water for 5 days, followed by a change to regular water. On day 7 after DSS treatment, *Il11* and *Egfp* mRNA expression in the colon was determined by qPCR. Results are mean ± SE (n = 9 mice). **b, c** Appearance of IL-11^+^ cells in submucosal tissues of the colon of *Il11-Egfp* reporter mice on post-DSS treatment day 5 (b) or day 1 (**c**). Colonic tissue sections from untreated or DSS-treated *Il11-Egfp* reporter mice were H&E stained or immunostained with anti-GFP antibody. Right panels show enlargements of the boxes (b). Scale bar, 100 μm. **d, e** Characterization of cell surface markers on IL-11^+^ cells. Colonic cells were prepared from the colon of *Il11-Egfp* reporter mice as in (**b**). We determined the percentages of EGFP^+^ (IL-11^+^) cells from the colon before and after DSS treatment (**d**). Cells were stained with the indicated antibodies, and marker expressions were analyzed on GFP-positive cells (**e**). Results are representative of three independent experiments. **f** Representative immunostaining of IL-11^+^ cells. Colonic tissue sections were prepared from *Il11-Egfp* reporter mice as in (**b**), and immunostained with the indicated antibodies and anti-GFP antibody. Results are merged images. Right panels are enlarged images from the boxes (n = 3–4 mice). White arrowheads indicate merged cells. **g** IL-11^+^ cells do not proliferate *in situ*. *Il11-Egfp* reporter mice were treated with DSS as in (**a**), and intraperitoneally administered BrdU (40 mg/kg) on day 6. On day 7, colonic sections were prepared and stained with anti-GFP and anti-BrdU antibodies. Results are representative images from three independent experiments. Scale bars, 100 μm, unless otherwise indicated. Statistical significance was determined by two-tailed unpaired Student’s *t*-test (**a**). *p < 0.05.

### IL-11^+^ Cells Express Genes Associated with Cell Proliferation and Tissue Repair

To further characterize the IL-11^+^ cells that appeared in colitis, we used a cell sorter to isolate IL-11^+^ cells as EGFP^+^ cells from the colon of DSS-treated *Il11-Egfp* reporter mice. These sorted EGFP^+^ cells were subjected to transcriptome analysis, and their gene expression profiles were compared with those of EGFP^−^ cells. Heat-map analysis revealed different gene expression patterns in these two populations (Fig. 4a). A volcano plot showed significantly elevated expressions of several genes, including *Il11*, *Mmp3*, and *Timp1*, in IL-11^+^ cells compared to IL-11^−^ cells (Fig. 4b). Gene Ontology (GO) enrichment analysis revealed that EGFP^+^ cells exhibited enriched expressions of genes associated with cell proliferation, organ morphogenesis, angiogenesis, and wound healing (Fig. 4c). qPCR confirmed that IL-11^+^ cells showed elevated levels of cytokines (*Il6* and *Il11*), chemokines (*Ccl2* and *Ccl11*), and genes associated with organ development (*Hgf* and *Tnfsf11*) (Fig. 4d). We also detected elevated expression levels of genes associated with colorectal cancer susceptibility loci (e.g., *Grem, Bmp2*, and *Bmp4*) and tumor development and invasion (e.g., *Wnt5a, Ereg*, *Mmp3*, *Mmp13,* and *Timp1*) (Figure 4D).

**Fig. 4.**
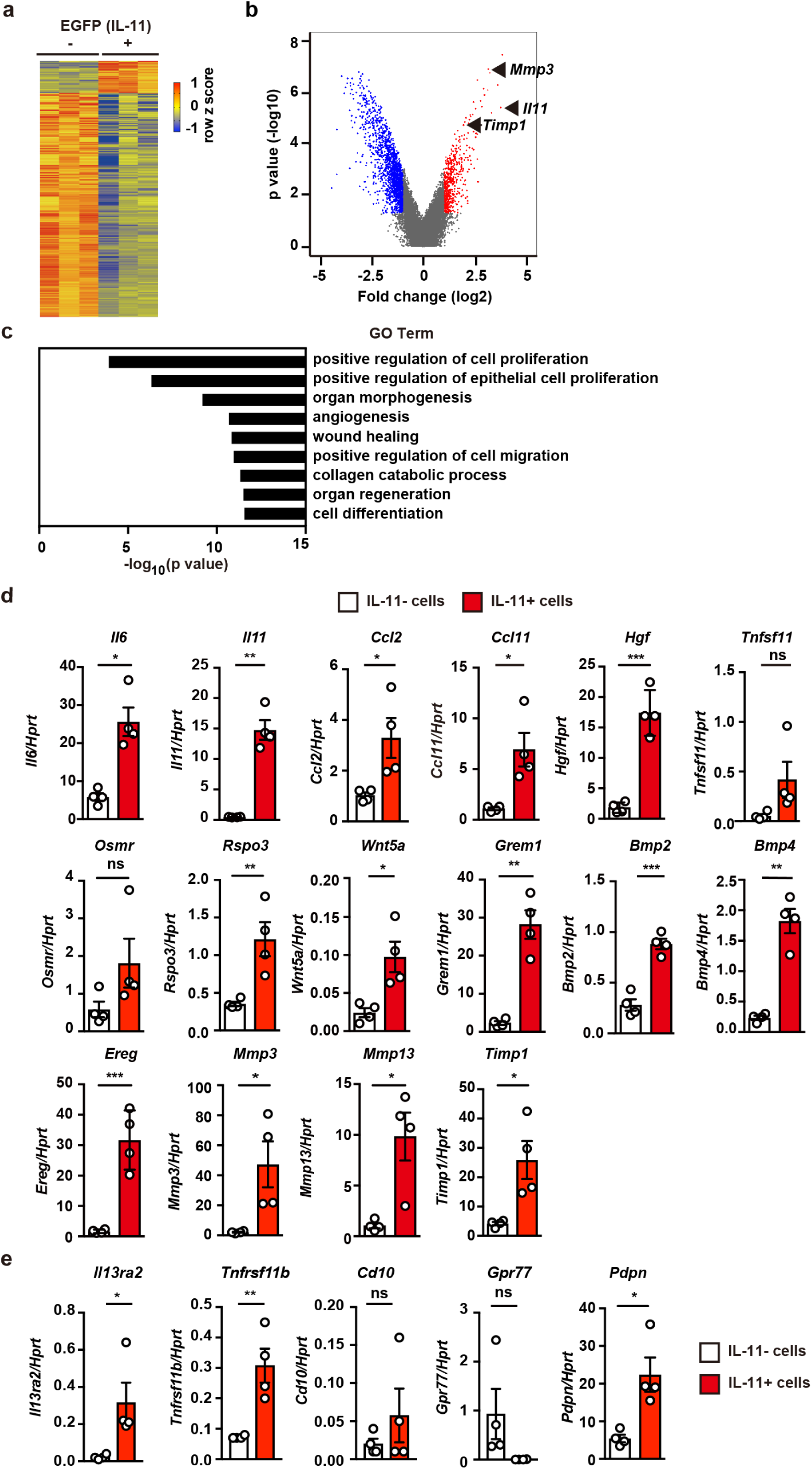
IL-11^+^ Cells Express Genes Associated with Cell Proliferation and Tissue Repair. On day 7 after DSS treatment, cells were isolated from the colon of *Il11-Egfp* reporter mice, and IL-11^+^ cells (EGFP^+^) were sorted by flow cytometry. We isolated mRNA from IL-11*^−^* and IL-11^+^ cells, and analyzed the gene expression by microarray analysis (n = 3). **a** Heat map of microarray gene expression of IL-11*^−^* and IL-11^+^ cells. Legend on right shows gene-expression color normalized by Z-score transformation. **b** Volcano plot of whole genes. Horizontal line indicates genes differentially regulated in IL-11^+^ cells compared to IL-11*^−^* cells, shown in log2. Vertical line indicates p values of statistical significance, shown in −log10. Significantly upregulated and downregulated genes are indicated by red and blue dots, respectively. Several upregulated genes are plotted. **c** Gene Ontology (GO) terms that were significantly enriched in IL-11^+^ cells compared to IL-11*^−^* cells. **d, e** Gene expressions were analyzed by qPCR. Results are mean ± SE (n = 4 mice). Statistical significance was determined by two-tailed unpaired Student’s *t*-test (**c, d**). *p < 0.05; **p < 0.01; ***p < 0.001; ns, not significant.

Of note, two unique subsets of fibroblasts termed inflammation-associated fibroblasts (IAFs)^9^ and cancer-associated fibroblasts (CAFs) with mediating chemotherapy resistance^10^ have been recently reported in ulcerative colitis (UC) patients and chemotherapy-resistant cancer patients, respectively. To investigate the relationship between IL-11^+^ fibroblasts and IAFs or CAFs with mediating chemotherapy resistance, we examined their hallmark gene expressions. We found elevated expressions of *Il13ra2* and *Tnfsfr11b* (markers of IAFs), but not *Cd10* or *Gpr77* (markers of CAFs with mediating chemotherapy resistance) in IL-11^+^ cells compared to IL-11*^−^* cells (Fig. 4e). These findings indicated that the IL-11^+^ cells appearing in the colon of DSS-treated mice were phenotypically similar to the IAFs observed in UC patients, but not CD10^+^GPR77^+^ CAFs. Thus, these cells were referred to as IL-11^+^ IAFs.

We next examined whether the gene expression profiles were similar between IL-11^+^ IAFs and IL-11^+^ CAFs. The genes enriched in IL-11^+^ IAFs, including *Ccl2, Osmr, Wnt5a, Ereg, Mmp13*, *Timp1*, and *Tnfrsf11b,* were also elevated in AOM/DSS-induced tumor tissues (Supplementary Fig. 4). Although we did not examine the gene expression profiles of IL-11^+^ CAFs from tumor tissues, our findings suggest that IL-11^+^ IAFs may be phenotypically similar to IL-11^+^ CAFs.

### The MEK/ERK Pathway is Involved in *Il11* Upregulation in Tumor Tissues

Since IL-11^+^ CAFs might promote tumor development, it is crucial to investigate the mechanisms by which inflammation or tumor cells induce IL-11 expression. We previously reported that oxidative stress induces *Il11* mRNA expression in an ERK/Fra-1-dependent manner^22, 23^. Thus, we tested whether DSS treatment induced oxidative stress in the colon. We observed enhanced oxidative stress in colonic cells after DSS treatment, and found that an antioxidant, N-acetyl cysteine (NAC) administration attenuated this DSS-induced oxidative stress (Fig. 5a). Additionally, DSS-induced oxidative stress was ameliorated by administration of antibiotics (Abx) (Fig. 5b), suggesting that bacterial infection induces oxidative stress. Consistent with these findings, both an antioxidant, NAC and Abx blocked the DSS-induced upregulation of *Il11* mRNA (Fig. 5c, d). Furthermore, DSS induced ERK phosphorylation, which was blocked by administration of the MEK inhibitor trametinib or NAC, accompanied by downregulation of *Il11* expression (Fig. 5e–g). These results indicated that oxidative stress-dependent ERK activation might contribute to *Il11* expression in the colon. Although several studies report that TGFβ induces IL-11 production in various cells^20, 21^, we found that neutralization of TGFβ signals using anti-TGFβ antibody did not block *Il11* expression in the colon after DSS treatment (Fig. 5h, i).

**Fig. 5.**
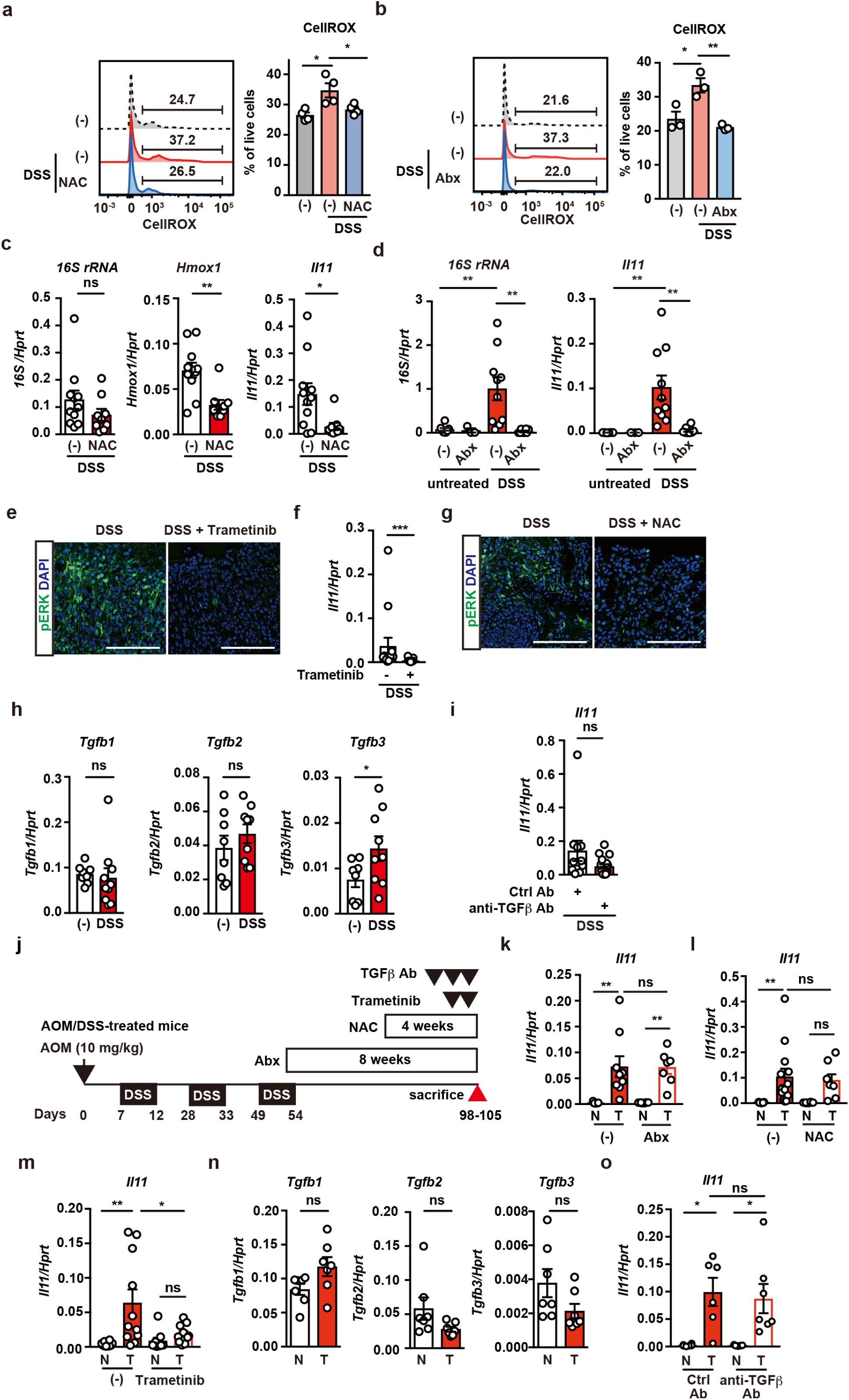
The MEK/ERK Pathway is Involved in *Il11* Upregulation in Tumor Tissues. **a, b** Wild-type mice were treated with DSS with or without NAC (**a**) or Abx (**b**). Colonic cells were prepared and stained with CellRox-green, and ROS accumulation was analyzed by flow cytometry. Left panels show representative histograms of ROS levels in colonic cells. Right panels show percentages of CellRox-green-positive cells of an individual mouse. Results are mean ± SE (n = 3–4 mice). **c** Wild-type mice were treated as in A. Colonic expressions of *16S rRNA, Hmox1,* and *Il11* mRNA were determined by qPCR. Results are mean ± SE (n = 11). **d** Abx blocked *Il11* mRNA upregulation in the colon of DSS-treated mice. Wild-type mice were untreated or treated with DSS in the absence or presence of Abx. On day 7 after DSS treatment, qPCR was performed to determine the expression of bacterial *16S rRNA* and *Il11* mRNA in the colon. Results are mean ± SE (n = 10). **e, f** Trametinib inhibits ERK phosphorylation and *Il11* mRNA expression in the colon of DSS-treated mice. Wild-type mice were treated with DSS in the absence or presence of trametinib, and then colonic sections were prepared and stained with anti-pERK antibody (**e**). Scale bars, 100 μm. *Il11* mRNA expression was determined by qPCR (**f**). Results are mean ± SE (n = 13–14). **g** NAC inhibits ERK phosphorylation in the colon of DSS-treated mice. Wild-type mice were treated with DSS in the absence or presence of NAC in the drinking water for 5 days. Colonic sections were stained with anti-pERK antibody. Results are representative of three independent experiments. Scale bar, 100 μm. **h** *Tgfbs* expression in the colon of DSS-treated wild-type mice. Wild-type mice were treated with DSS as in Fig. 4a. On day 7 after DSS treatment, qPCR was performed to determine *Tgfb1, Tgfb2*, and *Tgfb3* mRNA expression in the colon. Results are mean ± SE (n = 8–10 mice). **i** Treatment with anti-TGFβ antibody does not downregulate *Il11* mRNA expression in the colon of DSS-treated mice. Mice were intraperitoneally administered 100 μg anti-TGFβ antibody on days 2 and 4 after DSS administration, and sacrificed on day 5. *Il11* mRNA expression was determined by qPCR. Results are mean ± SE (n = 11–16 mice). **j** Schema of administration of various inhibitors in AOM/DSS-treated mice. Mice were treated with Abx for 8 weeks, NAC for 4 weeks, trametinib at −6 and −30 hours, or anti-TGFβ antibody on day −1, −3, −5 (just before sacrifice). **k–m, o** Mice were treated as in Figure 1A, and then treated with Abx (**k**) (n = 7–9 mice), NAC (**l**) (n = 7–11 mice), trametinib (**m**) (n = 10–12 mice), or anti-TGFβ antibody (n = 6–7 mice) (**o**) as in (**j**). On day 98–105 after AOM injection, mRNA was extracted from tumor and non-tumor tissues, and *Il11* expression was determined by qPCR. Results are mean ± SE. **n** Mice were treated as in Fig. 1a. *Tgfb1*, *Tgfb2,* and *Tgfb3* expressions in tumor and nontumor tissues were determined by qPCR. Results are mean ± SE (n = 7 mice). Statistical significance was determined using the unpaired two-tailed Student’s *t*-test (**c, h**), Mann-Whitney U test (**f, i**), two-way ANOVA with Bonferroni’s test (**d, k, l, m, o**), or one-way ANOVA with Tukey’s post-hoc test (**a, b**). *p < 0.05; **p < 0.01; ***p < 0.001; ns, not significant.

We next investigated whether *Il11* expression was induced in the tumor microenvironment in a manner similar to those in DSS-induced colitis. Administration of trametinib, but not Abx, NAC, or neutralizing antibody against TGFβ, decreased *Il11* expression in CAC after AOM/DSS treatment (Fig. 5j–o). Moreover, we observed a similar inhibitory effect of Trametinib, but not Abx, NAC, or neutralizing antibody against TGFβ in colon tumors in *Apc^min/+^* mice (Supplementary Fig. 5a–f). Together, these results suggested that *Il11* expression was induced in a MEK/ERK-dependent manner, although the upstream signals that induce activation of the MEK/ERK pathway could differ between cases of inflammation versus tumors. Of note, induction of *Il11* in the colon might be independent of TGFβ in the tumor microenvironment at least under our experimental conditions.

### IL-11 Preferentially Induces Signals to Fibroblasts

Previous studies report that IL-11 induces signals in colonic epithelial cells and fibroblasts^17, 29^. To determine which cells were the primary targets of IL-11, we examined the *Il11ra1* expression on colonic fibroblasts and normal colonic epithelial organoids. Since IL-22 induces signals in intestinal epithelial cells^36^, we also examined the IL-22 receptor expression on both cell types. Colonic epithelial organoids showed high expression of *Il22r*, but not *Il11ra1*, while colonic fibroblasts exhibited high expression of *Il11ra1*, but not *Il22ra1* (Fig. 6a). Consistently, robust STAT3 phosphorylation in colonic fibroblasts was induced by IL-11, but not IL-22 (Fig. 6b). To further characterize IL-11-inducible genes *in vivo*, we injected IL-11R agonist into wild-type mice^22^, which induced upregulation of many genes in the colon (Fig. 6c–e), some of which were enriched in IL-11^+^ IAFs (Fig. 6e). We performed qPCR to verify the induction of these genes (including *Hgf, Osmr, Rspo3*, *Mmp3, Timp1,* and *Pdpn*) in the colon after injection of IL-11R agonist (Supplementary Fig. 6). It is likely that IL-11 released from colonic fibroblasts might activate IL-11^+^ IAFs or CAFs in an autocrine or paracrine manner. Overall, our findings suggested that expression of IL-11-stimulated genes in IL-11^+^ CAFs might further modulate the tumor microenvironment.

**Fig. 6.**
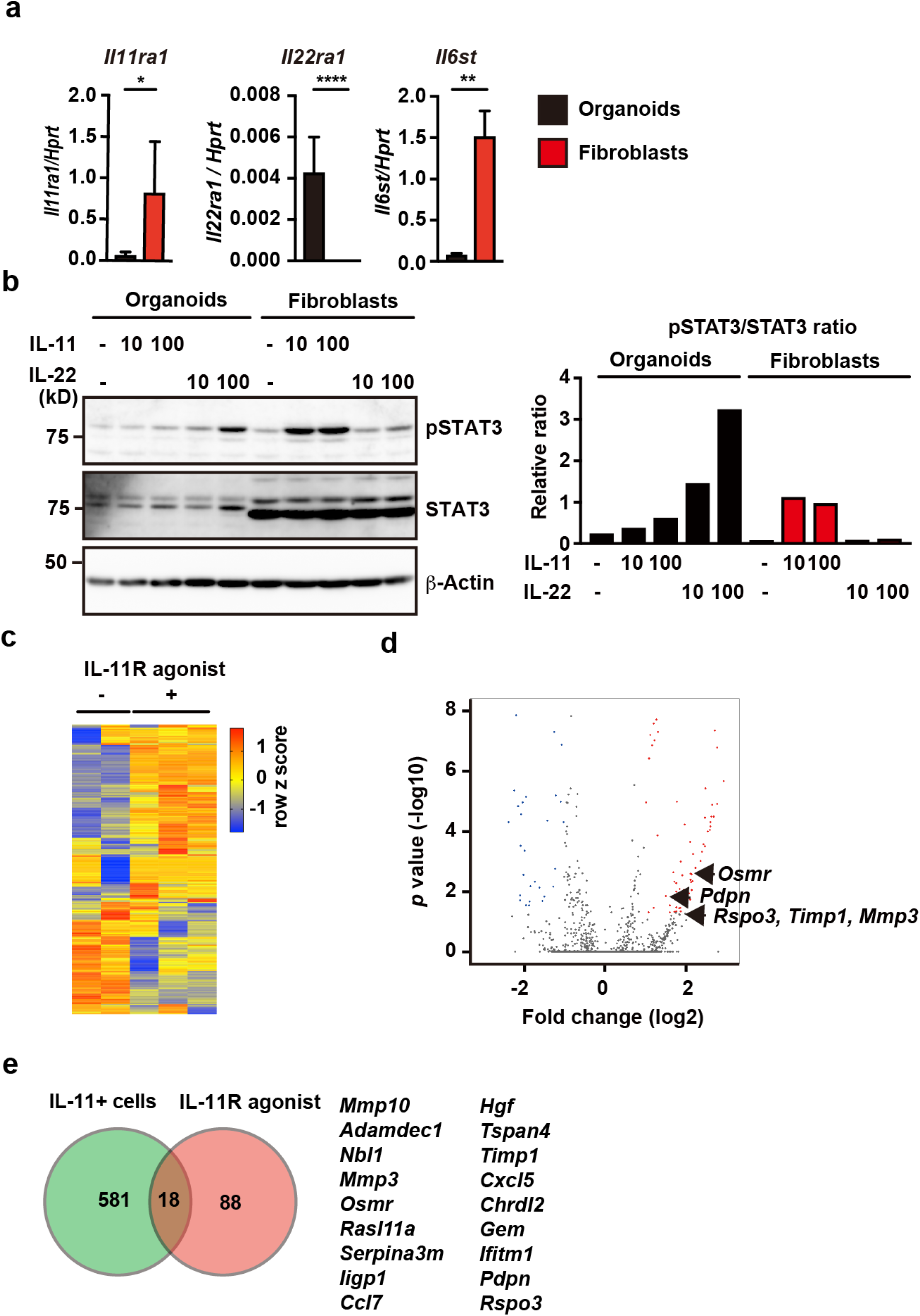
IL-11 Preferentially Induces Signals to Fibroblasts. **a** Relative expressions of *Il11ra1*, *Il22ra1*, and *Il6st* in colonic epithelial organoids and colonic fibroblasts. Colonic epithelial organoids and fibroblasts were established from wild-type mice as described in the methods. Expressions of the indicated genes were determined by qPCR. Results are mean ± sd of triplicate samples and representative of two independent experiments. **b** Colonic epithelial organoids and fibroblasts were unstimulated or stimulated for 30 min with IL-11 (10 or 100 ng/mL) or IL-22 (10 or 100 ng/mL). Total STAT3 and phosphorylated STAT3 (pSTAT3) were analyzed by Western blotting. STAT3 and pSTAT3 signaling intensities were calculated by Fiji, and the relative ratio of pSTAT3/STAT3 is shown. Results are representative of two independent experiments. **c** Administration of IL-11R agonist induces expression of the genes expressed in IL-11^+^ cells. We injected 8-week-old wild-type mice with 10 μg IL-11R agonist. At 3 hours after injection, mRNA was isolated from the colon, and gene expressions were analyzed by microarray analysis (n = 2 for untreated samples; n = 3 for injected samples). Heat map of microarray gene expression in the colon of untreated and treated mice (**c**). Legend on the right shows gene-expression color normalized by Z-score transformation. **d** Volcano plot of whole genes. Horizontal line indicates differentially regulated genes in the colon after IL-11R agonist injection compared to before injection, shown in log2. Vertical line indicates p values of statistical significance, shown in −log10. Significantly upregulated genes are indicated by red dots. Several upregulated genes are plotted. **e** Venn diagram of genes elevated in IL-11^+^ IAFs compared to IL-11*^−^* cells, and genes elevated in the colon with IL-11R agonist treatment compared to untreated colon. Overlapping area includes 18 genes. Statistical significance was determined by two-tailed unpaired Student’s *t*-test (**a**). *p < 0.05; **p < 0.01; ****p < 0.0001.

### Genes Enriched in IL-11^+^ IAFs Show Elevated Expression in Human Colorectal

#### Cancer

Assuming that the appearance of IL-11^+^ CAFs was correlated with CAC development in mice (Fig. 1, 2), we hypothesized that the genes enriched in IL-11^+^ IAFs might also be elevated in human colorectal cancer. We focused on genes with over two-fold greater expression in IL-11^+^ IAFs compared to IL-11^−^ cells (Fig. 4), and extracted 17 genes matching our criteria from the human cancer database (GSE33133). Intriguingly, subsets of genes elevated in IL-11^+^ IAFs, including *HGF, TNFSF11, MMP3, MMP13,* and *TIMP1*, were significantly upregulated in colon cancer tissues compared to normal mucosa (Supplementary Fig. 7a).

Since many of the genes enriched in IL-11^+^ IAFs were also upregulated in tumor tissues (Fig. 4d, e, Supplementary Fig. 7a), we hypothesized that the genes elevated in IL-11^+^ IAFs, along with *IL11*, might critically affect the prognosis of cancer patients. Thus, we compared disease-free survival of colorectal cancer patients according to the expression levels of genes enriched in IL-11^+^ IAFs. We found that reduced disease-free survival time was associated with higher expression of *IL6*, *MMP13*, and *TIMP1*, but not of *IL6/IL6R*, *IL11*, or *IL11/IL11R* (Supplementary Fig. 7b). Notably, higher expression of *IL6/IL6R/IL11/IL11R/GP130* was associated with significantly diminished disease-free survival time. These results suggested that prognosis of human colorectal cancer patients may be critically determined by IL-11^+^ IAFs and, possibly, by IL-11^+^ CAFs, but not by IL-11.

#### IL-11 Expression is Correlated with Progression of Human Cancers

To characterize IL-11^+^ cells in human colon tumor samples, we performed staining with anti-human IL-11 antibody. First, we verified that anti-human IL-11 antibody detected endogenous IL-11, using lysates of the human breast cancer cell line MDA-MB-231, in the absence or presence of siRNA against human *Il11* (Fig. 7a). IL-11^+^ cells were scarcely detected in normal colonic tissues, but were numerous in adenomas and in early and advanced colorectal cancer tissues (Fig. 7b). Intriguingly, the relative area of IL-11^+^ cells and the IL-11 signaling intensity were increased in advanced colon cancers compared to normal tissues (Fig. 7c, d). Moreover, positive staining for IL-11 was observed in vimentin-positive stromal cells, E-cadherin-positive tumor cells, and CD45-positive hematopoietic cells (Fig. 7e). Thus, IL-11^+^ cells comprised heterogeneous cell populations in the tumor microenvironment, although the majority were fibroblasts.

**Fig. 7.**
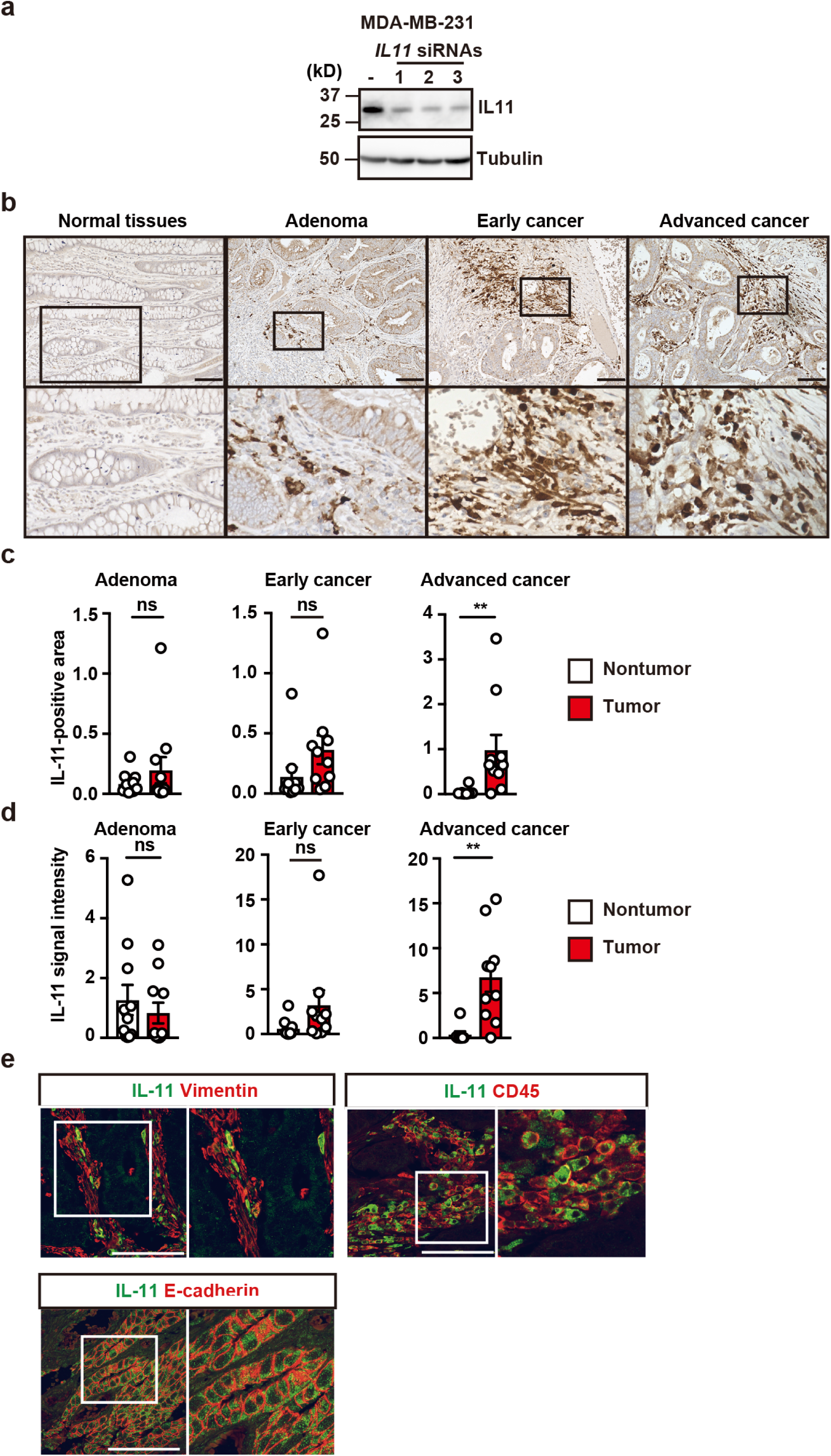
IL-11 Expression is Correlated with Progression of Human Cancers. **a** Specificity of anti-IL-11 antibody used in the study. MDA-MB-231 cells were treated with control or three different siRNAs against human *Il11*, and IL-11 expression in cell lysates was analyzed by Western blotting with anti-IL-11 and anti-tubulin antibodies. Results are representative of two independent experiments. **b-d** Adenomas (n = 10), early (n = 10) and advanced (n = 10) colorectal cancers were stained with anti-IL-11 antibody (**b**). Representative staining of respective tumors and adjacent normal tissues. Scale bars, 100 μm. Areas (**c**) and intensities (**d**) of IL-11^+^ signals were calculated as in the methods. Results are mean ± SE. **e** Tissues of advanced colorectal cancers were stained with anti-vimentin, anti-CD45, or anti-E-cadherin antibodies along with anti-IL-11 antibody (n = 3). Right panels are enlarged images of the white box in the left panels. Signaling intensities stained with the indicated antibodies on the white arrows were calculated and plotted in the right panels. Scale bars, 100 μm. Statistical significance was determined by Mann-Whitney U test (**c, d**). **p < 0.01; ns, not significant.

## DISCUSSION

In the present study, we generated *Il11-Egfp* reporter mice to characterize IL-11^+^ CAFs or IL-11^+^ IAFs in different murine tumor and colitis models. We found that IL-11^+^ cells in tumor tissues constituted heterogenous cell populations, including stromal, epithelial, and hematopoietic cells, and that *Il11ra^−^*^/*−*^ or *Illl^−^*^/*−*^ deletion attenuated CAC development in mice. Reciprocal BM transfer experiments revealed that stromal cells critically contributed to tumor progression; thus, we focused on IL-11^+^ CAFs. IL-11 activated IL-11^+^ colonic fibroblasts. Transcriptome analysis showed that IL-11^+^ IAFs expressed genes associated with tissue repair and cell proliferation. Moreover, some of the genes enriched in IL-11^+^ IAFs were elevated in human colorectal cancer tissues, and their high expression was associated with poor prognosis of human colorectal cancer. Thus, our results suggest that a feed-forward loop between IL-11 and IL-11^+^ CAFs may contribute to tumor development.

The newly developed *Il11-Egfp* reporter mice enabled us to characterize the IL-11^+^ IAFs and CAFs that appeared in mouse models of colitis and CAC, respectively. In the colitis model, IL-11^+^ IAFs exclusively expressed stromal cell markers, such as Thy1 and podoplanin. A recent cRNA-seq analysis of colon biopsies from healthy individual and UC patients revealed that colon samples from UC patients exhibited a unique subset of fibroblasts, termed IAFs, that express *IL11*, *IL24*, *IL13RA2,* and *TNFSFR11B*^9^. Here we found elevated expression of these genes in IL-11^+^ IAFs compared to IL-11*^−^* cells. Thus, IL-11 is a hallmark of IAFs. Since IL-11^+^ IAFs expressed genes associated with cell proliferation and tissue repair, IL-11^+^ IAFs might contribute to colitis attenuation in human UC patients.

In contrast to in DSS-induced colitis, in tumor tissues, we detected numerous IL-11^+^ cells expressing epithelial or hematopoietic cell markers, including EpCAM or CD45.2. Consistently, previous studies report that IL-11^+^ cells are derived from stromal, epithelial, or hematopoietic cells^17, 24^. Our present results revealed that IL-11^+^, EpCAM^+^, and E-cadherin^+^ epithelial cells might be tumor cells themselves, whereas we did not fully investigate the nature and origin of CD45.2^+^ cells. Future scRNA-seq analysis of IL-11^+^ cells will further elucidate the origin and characterization of heterogenous populations of IL-11^+^ cells in the tumor microenvironment.

Tumors cells may support tumor growth by educating the surrounding fibroblasts, which are referred to as CAFs^8, 37, 38^. On the other hand, a recent study reported that surface expression of CD10 and GPR77 may be markers of CAFs that mediate chemotherapy resistance in human cancer through IL-6 and IL-8 production ^10^. CD10 and GPR77 expression were not elevated in IL-11^+^ IAFs or CAC in mice, suggesting that IL-11^+^ IAFs and possibly IL-11^+^ CAFs might be different from human CD10^+^GPR77^+^ CAFs.

Elucidation of the mechanisms underlying IL-11 production by stromal cells in the tumor microenvironment may be crucial for understanding how tumor cells instruct stromal cells. Previous studies demonstrate that TGFβ induces IL-11 upregulation in various cell types^20, 21^. Indeed, we found that TGFβ stimulation induced IL-11 production by colonic fibroblasts. Moreover, in murine xenograft models, human tumor cell lines that ectopically express human TGFβ1 can elicit IL-11^+^ CAFs^24^. However, in our present study, anti-TGFβ antibody treatment did not downregulate *Il11* expression in colonic adenomas of *Apc^min/+^* mice, or in AOM/DSS-induced CAC. Furthermore, culture supernatants of tumor organoids did not induce upregulated *Il11* expression by colonic fibroblasts in the absence or presence of anti-TGFβ antibody. Overall, although TGFβ *per se* was able to induce IL-11 production, TGFβ may not be involved in the induction of IL-11^+^ CAFs in the tumor microenvironment, at least under our experimental conditions. On the other hand, blockade of the ERK pathway attenuated *Il11* expression in the colon of mice after DSS treatment, in colon adenomas of *Apc^min/+^* mice, and in AOM/DSS-induced CAC in mice. Thus, the signaling pathway(s) leading to MEK/ERK activation might be critically involved in IL-11 production. Further investigations are needed to address this subject.

Previous studies show that IL-11 and IL-22 induce proliferation and cell survival of colonic epithelial cells through STAT3 activation^15, 36^. Indeed, under our experimental conditions in colonic epithelial organoids, IL-22 induced strong STAT3 phosphorylation, whereas IL-11 induced only weak STAT3 phosphorylation. In sharp contrast, IL-11 induced robust phosphorylation of STAT3 in colonic fibroblasts. Thus, it appears that colonic fibroblasts produced IL-11 in response to factors from tumor cells, which, in turn, activated and induced STAT3 phosphorylation in colonic fibroblasts in an autocrine or paracrine manner. This feed-forward loop between IL-11 and IL-11^+^ CAFs might contribute to tumor development.

CAC development was attenuated in *Il11ra1^−^*/*^−^* and *Il11^−^*/*^−^* mice treated with AOM/DSS, but deletion of these genes did not dramatically affect tumor development compared to in a previous study^17^. The reason for these discrepancies is currently unknown. However, it is possible that the genetic background of the mice (C57/BL6 vs. 129/C57/BL6 mixed background) might affect the tumor cells’ dependence on IL-11 signaling. On the other hand, our transcriptome analysis revealed that IL-11^+^ CAFs expressed various genes associated with tissue repair and cell proliferation. Indeed, we found that high expression of the genes enriched in IL-11^+^ IAFs (including *IL6*, *MMP3*, and *TIMP1*, but not *IL11* itself) was associated with reduced disease-free survival of CRC patients. Thus, it would be interesting to investigate whether depletion of IL-11^+^ CAFs might have profound effects on colorectal cancer progression.

## METHODS

### Reagents

Recombinant mouse IL-11 (R&D), IL-22 (Biolegend), mouse TGFβ1 (Biolegend), N-acetyl cysteine (NAC) (Nakalai), and Trametinib (LC Laboratories) were purchased from the indicated sources. The following antibodies used in this study were obtained from the indicated sources: anti-phospho-ERK (4370, CST), anti-Ki67 (ab16667, Abcam), anti-GFP (GFP-Go-Af1480 or GFP-Rb-Af2020, Frontier Institute), anti-BrdU (BU1/75, BIO-RAD), anti-IL-11 (LS-C408373, LSBio), anti-CD45 (13917, CST), anti-CD45 (IR751, Dako), anti-podoplanin (127403, BioLegend), anti-α-SMA (ab5694, Abcam), anti-collagen I (ab34710, Abcam), anti-collagen IV (ab6586, Abcam), anti-E-cadherin (560062, BD Biosciences), anti-E-cadherin (NCH-38, Dako), anti-vimentin (9856, CST), anti-phospho-STAT3 (9145, CST), anti-STAT3 (SC-482, Santa Cruz), anti-β-Actin (622102, Biolegend), and anti-tubulin (T5168, Sigma-Aldrich). Anti-horseradish peroxidase (HRP) -conjugated anti-rabbit IgG (NA934), HRP-conjugated anti-rat IgG (NA935), and HRP-conjugated anti-mouse IgG (NA931) antibodies were from GE Healthcare. Alexa Fluor 488-conjugated donkey anti-rabbit IgG (A21206), Alexa Fluor 594-conjugated donkey anti-rabbit IgG (A21207), Alexa Fluor Plus 594-conjugated donkey anti-rabbit IgG (A32754), Alexa Fluor 647-conjugated donkey anti-rabbit IgG (A31573), Alexa Fluor 594-conjugated donkey anti-mouse IgG (A21203), and Alexa Fluor 488-conjugated donkey anti-goat IgG (A11055) antibodies, and Alexa Fluor 594-conjugated streptavidin (S11227) were purchased from Invitrogen.

Unless otherwise indicated, the following antibodies used for flow cytometry were obtained from TONBO Biosciences; anti-CD11b (20-0112, clone M1/70), anti-CD16/CD32-mAb (2.4G2) (made in house), anti-CD24a (Biolegend, 101813, clone M1/69), anti-CD31 (eBioscience, 17-0311-82, clone 390), anti-CD34 (eBioscience, 13-0341-81, clone RAM34), anti-CD45.1 (35-0453, clone A20), anti-CD45.2 (20-0454, clone 104), anti-EpCAM (BioLegend, 118214, clone G8.8), anti-Thy1.2 (20-0903, clone 30-H12), anti-podoplanin (BioLegend, 127414, clone 8.1.1), anti-TER-119 (BioLegend, 116212, clone TER-119), anti-MHC Class II (Miltenyi Biotec, 130-102-139, clone M5/114.15.2), anti-ICAM-1 (BD Biosciences, 561605, clone 3E2), anti-VCAM-1 (BioLegend, 105718, clone 429), anti-Sca-1 (BioLegend, 122512, clone E13161.7), anti-Lyve1 (eBioscience, 50-0443-80, clone ALY7), anti-PDGFRα (eBioscience, 17-1401-81, clone APA5), and Streptavidin APC (eBioscience).

The hybridoma cell line (1D11)^39^ that produces neutralizing antibody against all TGFβ isoforms (β1, β2, and β3) was purchased from ATCC, and anti-TGFβ antibody was produced in house. Control mouse IgG was purchased from Sigma-Aldrich (I5381).

### Mice

*Il11^-/-^* mice (generated in our lab) and *Il11ra1*^-/-^ mice (provided by L. Robb) were previously described^34 40^. *Apc^min/+^* (002020) were purchased from Jackson Lab. C57BL/6 (CD45.2^+^) and C57BL/6-SJL (CD45.1^+^) mice were purchased from Japan-SLC.

Mice in different cages or derived from different sources were cohoused for 2 weeks for normalization of the microbiota composition before experiments. All animals were housed and maintained under specific pathogen-free conditions in the animal facility at Juntendo University School of Medicine or Toho University School of Medicine. All experiments were performed according to the guidelines approved by the Institutional Animal Experiment Committee of Juntendo University School of Medicine or Toho University School of Medicine (19-51-414, 19-51-411).

### Generation of *Il11-Egfp* reporter mice

The *Egfp* reporter gene was introduced into the BAC clone (RP23-285B12) by two-step Red/ET recombineering technology according to the manufacturer’s protocol (Gene Bridges). In the first step, a *rpsL-neo* cassette included in the kit was amplified by PCR using a primer set (*Il11*-ET1-F2: ACTCCCTCAGACCCAGAGTTTGGCCTGATTTCTCCCTTCTGTCCACAGGTGGCCTGGTGATGATGGCGGGATCG and *Il11*-ET1-R2: ACGACTCTATCTGGCCAGAGGCTCAGCACCACCAGGACCAGGCGACAAACTCAGAAGAACTCGTCAAGAAGGCG) and inserted into the target region of the BAC clone. In a second step, the *rpsL-neo* cassette in the modified BAC clone was replaced with *Egfp-polyA* cassette which was amplified from *Egfp*-expression vector (pAWZ) using a primer set (*Il11*-ET2-F2: ACTCCCTCAGACCCAGAGTTTGGCCTGATTTCTCCCTTCTGTCCACAGGTATGGTGAGCAAGGGCGAG and *Il11*-ET2-R2: ACGACTCTATCTGGCCAGAGGCTCAGCACCACCAGGACCAGGCGACAAACctCTAGTGGATCATTAACGCTTAC). A resultant clone designated as *IL11-Egfp* was verified by restriction digestion of BAC DNA and by sequencing.

Intracytoplasmic sperm injection (ICSI) was performed as previously described with slightly modifications^41^. The mixture of sperm and *Il11-Egfp* DNA was diluted with Hepes-modified CZB containing 12% polyvinylpyrrolidone (Sigma-Aldrich) before being used for ICSI. Injections were performed by micromanipulators (Leica) with a PMM-150 FU piezo-impact drive unit (Prime Tech) using a blunt-ended mercury-containing injection pipette. After discarding the midpiece and tail, the head of spermatozoa was injected into an oocyte from C57/BL6 mice. Oocytes matured into two-cell stage embryos at 24 hours after injection, and then two-cell stage embryos were transferred to oviducts of pseudopregnant females. Transgenic founders were backcrossed to C57/BL6J mice for several generations. Among them, one line exhibited an intimate correlation of *Il11* and *Egfp* expressions were selected and used for further experiments.

### Induction of DSS-induced colitis and colitis-associated cancer (CAC) in mice

Nine- to fifteen-week-old wild-type or *Il11-Egfp* reporter mice received 1.5 % DSS (MW: 36,000-50,000 D; MP Biomedicals) *ad libitum* in drinking water for 5 days, which then was changed to regular water. To reduce gut commensal microflora, mice received mixtures of antibiotics in drinking water containing ampicillin (1 g/L, Sigma-Aldrich), kanamycin (0.4 g/L, Sigma-Aldrich), gentamicin (0.035 g/L, Sigma-Aldrich), metronidazole (0.215 g/L, Sigma-Aldrich), vancomycin (0.18 g/L, Sigma-Aldrich), and colistin (0.042 g/L, Sigma-Aldrich). Administration of antibiotics into mice started at 4 weeks before DSS treatment and continued during DSS treatment. To attenuate oxidative stress in DSS-treated mice, mice received NAC (10 g/L) along with DSS in drinking water for 5 days. To neutralize TGFβ in DSS-treated mice, mice were intraperitoneally injected with anti-TGFβ antibody (1D11) or control mouse IgGs (5 mg/kg) on day 2 and day 4 after DSS treatment. To inhibit ERK activation in DSS-treated mice, a MEK inhibitor, Trametinib (2 mg/kg) was administered into mice by gavage at 30 and 6 hours before the start of DSS treatment at a fine suspension in 0.5% Hydroxypropyl Cellulose (Alfa Aesar) and 0.2 % Tween-80.

To induced CAC, 9- to 15-week-old mice were intraperitoneally injected with 10 mg/kg AOM (Sigma-Aldrich). One week later, mice received 1.5 % DSS *ad libitum* in drinking water for 5 days, followed by 2 weeks of regular water, and this was repeated for two additional cycles. To reduce numbers of commensal bacteria and oxidative stress in AOM/DSS-treated mice or *Apc^min/+^* mice, we administered mixtures of antibiotics and NAC into the indicated mice for the last 8 weeks and 4 weeks just before sacrifice, respectively. Trametinib (2mg/kg) (−6 and −30 hours) or anti-TGFβ antibody (5 mg/kg) (on day −1, −3, −5) were administered into AOM/DSS-treated mice or *Apc^min/+^* mice at the indicated days just before sacrifice.

### Flow cytometry

To isolate IL-11^+^ cells, the colon was removed from DSS-treated *Il11-Egfp* reporter mice. Then, the colon was minced with scissors and digested in RPMI 1640 containing 100 U/mL Penicillin and 100 μg/mL Streptomycin, 1mg/mL Collagenase (Wako), 50 μg/mL DNase (Roche), 0.5 mg/mL Dispase (Roche), and 2% (v/v) fetal bovine serum (FBS, Gibco) for 60 min. Single cell suspensions were prepared, and cells were stained with the indicated antibodies and analyzed by LSRFortessa X-20 (BD Biosciences) or BD Verse (BD Biosciences). Data were processed by FlowJo software (FlowJo). To further characterize IL-11^+^ cells, EGFP^+^ cells were sorted by MoFlo Astrios cell sorter (Beckman Coulter) and subjected to microarray analysis.

To isolate IL-11^+^ cells from tumors in the colon of *Il11-Egfp* reporter mice treated with AOM/DSS or *Apc^min/+^;Il11-Egfp* mice, tumor tissues were removed from non-tumor tissues. Then, single cell suspension from tumor tissues was prepared as described above. Cells were stained with the indicated antibodies and expression of various cell surface markers on GFP^+^ cells were analyzed by flow cytometry.

### Generation of bone marrow chimeras

After BM cells were prepared from *Il11^+/+^* or *Il11^-/-^* mice (CD45.2), 3-5 x 10^6^ BM cells were transferred to 8-week-old recipient mice [C57BL/6-SJL mice (CD45.1)] that had been exposed to lethal irradiation (9.0 Gray). In reciprocal BM transfer experiments, BM cells from wild-type C57BL/6-SJL mice (CD45.1) were transferred to *Il11^+/+^* or *Il11^-/-^* mice (CD45.2) that had been exposed to lethal irradiation (9.0 Gray). At 2 to 3 months after transfer, peripheral blood mononuclear cells were collected and stained with FITC-conjugated anti-CD45.1 and allophycocyanin (APC)-conjugated anti-CD45.2 antibodies. The chimerism of bone marrow cells was calculated by counting numbers of CD45.1^+^ and CD45.2^+^ by flow cytometry. Average chimerisms were more than 90 %.

### Quantitative PCR (qPCR) Assays

Total RNAs were extracted from the indicated tissues of mice by using TRI Reagent (Molecular Research Center) or Sepasol II Super (Nacalai Tesque), and cDNAs were synthesized with the RevertraAce qPCR RT Kit (Toyobo). To remove residual DSS, mRNAs prepared from the colon of mice treated with DSS were further purified by LiCl precipitation as described previously ^42^. Quantitative polymerase chain reaction (qPCR) analysis was performed with the 7500 Real-Time PCR detection system with CYBR green method of the target genes together with murine *Hprt* an internal control with 7500 SDS software (Thermo Fisher Scientific). The primers used in this study are shown in Table S1.

### Microarray analysis

We compared gene expression profiles of RNAs from EGFP-positive and negative cells from the colon of *Il11-Egfp* reporter mice. EGFP^+^ and EGFP^-^ cells were sorted from the colon of *Il11-Egfp* mice (n = 3 mice) on day 7 after DSS administration, total RNAs were extracted using TRI-reagent according to the manufacturer’s instructions (Molecular Research Center). Purified RNAs were subjected to a LiCl purification to remove residual DSS. cDNA was synthesized with the Ambion WT Expression Kit (Affymetrix) and labeled with GeneChip WT Terminal Labeling Kit (Affymetrix) according to the manufacturer’s protocol. Affymetrix GeneChip Hybridization wash stain kit was used to hybridize samples to GeneChip Mouse Gene ST 1.0 ST Array (Affymetrix). Signals were scanned with the Genechip Scanner 3000 7G, and obtained data were analyzed with Affymetrix Expression Console software (Affymetrix). The gene expression was analyzed using GeneSpring (Silicon Genetics). Functional enrichment analysis of differentially expressed genes from the data of microarray analysis was performed using DAVID Bioinformatics Resources^43, 44^. Data were deposited in NCBI as a GSE140411.

To identify target genes by IL-11 in the colon, we intravenously injected 10 μg of IL-11R agonist into wild-type mice (n=2-3 mice). Generation, purification, and characterization of IL-11R agonist were described previously^22^. Mice were sacrificed at 3 hours after injection. Total RNAs were extracted from the colon of mice by using Sepasol II Super. Labeled cRNA were prepared from total RNA using the Agilent’s Quick Amp Labeling Kit (Agilent). Following fragmentation, cRNA were hybridized to SurePrint G3 Mouse Gene Expression 8×60K (Agilent) according to the manufacturer’s instruction. Raw data were extracted using the software provided by Agilent Feature Extraction Software (v11.0.1.1). The raw data for same gene was then summarized automatically in Agilent feature extraction protocol to generate raw data text file, providing expression data for each gene probed on the array. Array probes that have Flag A in samples were filtered out. Selected processed signal value was transformed by logarithm and normalized by quantile method. Statistical significance of the expression data was determined using fold change and LPE test in which the null hypothesis was that no difference exists among 2 groups. Hierarchical cluster analysis was performed using complete linkage and Euclidean distance as a measure of similarity. Gene-Enrichment and Functional Annotation analysis for significant probe list was performed using Gene Ontology (http://geneontology.org). All data analysis and visualization of differentially expressed genes was conducted using R 3.0.2 (www.r-project.org). Data were deposited in NCBI as a GEO accession number GSE141643.

### Immunohistochemistry (IHC)

Tissues were fixed in 10% formalin and embedded in paraffin blocks. Paraffin-embedded colonic sections were used for H&E staining, immunohistochemical, and immunofluorescence analyses. For immunohistochemistry, paraffin-embedded sections were treated with Instant Citrate Buffer Solution (RM-102C, LSI Medicine) or Target Retrieval Solution (S1699, Dako) as appropriate to retrieve antigen. Then tissue sections were stained with the indicated antibodies, followed by visualization of Alexa-conjugated secondary antibodies, or biotin-conjugated secondary antibodies followed by Streptavidin-HRP (Vector Laboratories).

In human tumor samples, tissue sections were preincubated with MaxBlock^TM^ Autofluorescence Reducing Kit (MaxVision Biosciences) according to the manufacturer’s instructions. After blocking, tissue sections were stained with the indicated antibodies as described above.

Pictures were obtained with an all-in-one microscope (BZ-X700, Keyence) or BX-63 (Olympus) and analyzed with BZ-X Analyzer (Keyence) or cellSens (Olympus) software. Confocal microscopy was performed on an LSM 880 (Zeiss) or A1R (Nikon). Images were processed and analyzed using the ZEN software (Zeiss) or the NIS-Elements (Nikon).

### Histological scoring of severity of colitis

Severity of colitis was evaluated by the following criteria, including infiltration of inflammatory cells and the degree of tissue damage as previously described ^45^. Infiltration of inflammatory cells; 0, the presence of rare inflammatory cells in the colonic lamina propria; 1, increased numbers of inflammatory cells; 2, confluence of inflammatory cells; 3, extending of inflammatory cells into the submucosa and transmural extension of the inflammatory cell infiltrate. The degree of tissue damage; 0, absence of mucosal damage; 1, discrete focal lymphoepithelial lesions; 2, mucosal erosion and ulceration; 3, extensive mucosal damage and extension through deeper structure of the intestinal tract wall. Total severity scores ranging from 0 to 6 represent that 0 and 6 indicate no changes and extensive cell infiltration with tissue damage, respectively.

### Organoid Culture and transplantation of tumor organoids

Mouse colon organoids were established from isolated crypts of wild-type mice as previously described with slight modification^46^. Obtained colonic organoids were cultured in Advanced DMEM/F12 (Thermo Fisher Scientific) containing 10 mM HEPES (Thermo Fisher Scientific), 1 x GlutaMAX (Thermo Fisher Scientific), 1 x B27 (Thermo Fisher Scientific), N-acetyl cysteine (1 μM), murine epidermal growth factor (50 ng/mL, Peprotech), murine Noggin (100 ng/mL, Peprotech), 0.5 mM A83-01 (Tocris), 3 μM SB202190 (Sigma), 1 μM nicotinamide (Sigma), Afamin and Wnt3A-condition medium (CM) (final concentration of 50%) which was provided by J. Takagi^47^ and R-spondin 1-CM (final concentration of 10%) (Trevigen).

The small intestinal tumor organoids from *villin-CreER-Apc^min/+^-Kras^+/LSL-G12D^-Tgfbr2^flox/flox^-Tp53^+/LSL-R270H^* (AKTP) mice were described previously ^35^. AKTP organoids were cultured in Advanced DMEM/F12 containing 10 mM HEPES, 1 x GlutaMAX, 1 x B27, N-acetyl cysteine (1 μM), murine epidermal growth factor (50 ng/mL), murine Noggin (100 ng/mL).

For transplantation, AKTP organoids were mechanically dissociated, and 3 x 10^5^ organoid cells mixed in Matrigel were injected into the subepithelial tissues of the rectum of *Il11-Egfp* reporter mice. At five weeks after transplantation, tumors were removed and subjected to histological analysis.

### Isolation and stimulation of colonic fibroblasts

To isolate colonic fibroblasts, single cell suspensions were prepared as described above and cultured in DMEM containing 10% FBS, 1% GlutaMAX (Thermo Fisher Scientific), 10 mM HEPES, 1% MEM Non-Essential Amino Acids Solution (100x) (Nacalai Tesque), 100 U/mL Penicillin, 100 μg/mL Streptomycin, and 2.5 μg/mL of Amphotericin B (Sigma). Culture medium was changed every day until colonic fibroblasts started to grow spontaneously.

To detect phosphorylation of STAT3 in colonic fibroblasts, colonic fibroblasts were stimulated with the indicated concentrations of IL-11 or IL-22 for 30 min. Total and phosphorylated STAT3 were analyzed by Western blotting.

### Western Blotting

Cells were lysed in a RIPA buffer (50mM Tris-HCl, pH 8.0, 150mM NaCl, 1% Nonidet P-40, 0.5% deoxycholate, 0.1% SDS, 25 mM β-glycerophosphate, 1mM sodium orthovanadate, 1mM sodium fluoride, 1 mM PMSF, 1 μg/ml aprotinin, and 1 μg/ml leupeptin). After centrifugation, cell lysates were subjected to SDS-PAGE and transferred onto polyvinylidene difluoride membranes (Millipore). The membranes were immunoblotted with the indicated antibodies. The membranes were developed with Super Signal West Dura extended duration substrate (Thermo Scientific) and analyzed by Amersham Biosciences imager 600 (GE Healthcare). In some experiments, blots were quantified using freeware program Fiji.

### Enzyme-linked immunosorbent assay (ELISA)

Concentrations of murine IL-11 in the culture supernatants were determined by ELISA according to the manufacturers’ instruction (R&D Systems).

### Knockdown of *Il11* by siRNAs

A human breast cancer cell line, MDA-MB-231 cells were provided by T. Sakamoto. MDA-MB-231 cells were maintained in DMEM containing 10% FBS. MDA-MB-231 cells were transfected with control or *Il11* siRNAs using Lipofectamine 2000 (Invitrogen) at a final concentration of 50 nM. At 36 h after transfection, knockdown efficiency of IL-11 by siRNAs was analyzed by immunoblotting with anti-IL-11 antibody using cell lysates. Stealth RNAi^TM^ siRNAs Negative Control, Med GC and human *Il11,* NM_000641.3 (HSS179893, HSS179894, HSS179895) were purchased from Thermo Fisher Scientific.

### Comparison of the expression of enriched genes in IL-11^+^ IAFs between human normal mucosa and colon cancer tissues

The publicly available data set (GSE33113) was obtained from the Gene Expression Omnibus (GEO). This data set contains gene expression data from 90 colon cancer patients and 6 healthy persons. The expression data of enriched genes in IL-11^+^ IAFs was retrieved from the data set (GSE33113) and their signaling intensities of each gene in normal mucosa and colon cancer tissues were compared.

### Survival analysis

Two publicly available data sets (GSE17536 and GSE17537) were obtained from the Gene Expression Omnibus (GEO). These data sets contain gene expression data and disease-free survival information from 232 primary CRC patients. The expression levels of enriched genes in IL-11^+^ IAFs in these data sets was classified by the hierarchical clustering. Each enriched gene in IL-11^+^ IAFs was correlated with survival using the Kaplan-Meier method. Statistical significance was analyzed by Mantel-Cox log-rank test. R software was used for all statistical analysis.

### Human colorectal cancer tissues

Human colon tumors and adjacent normal tissues were obtained from Toho University Omori Medical Center and analyzed in accordance with approval by the Ethics Committee of Toho University School of Medicine (A16111). Colon tumors included 10 cases of adenomas, 10 cases of early cancers, and 10 cases of advanced cancers. Histological assessment of adenomas and adenocarcinomas was according to the guidelines of the World Health Organization ^48^. Early and advanced colon cancers were determined according to the TNM criteria (pT1, early cancer; pT2-4, advanced cancers)^49^.

### Statistical analysis

Statistical significance was determined by the unpaired two-tailed Student’s *t*-test, Mann-Whitney U test, two-way ANOVA with Boniferroni’s test, or one-way ANOVA with Tukey’s post-hoc test, Mantel-Cox log-rank test as indicated. *p < 0.05 was considered to be statistically significant. All statistical analysis was performed with Graph Pad Prism 7 software (GraphPad Software).

## ACKNOWLEDGEMENTS

We thank T. Sakamoto for MDA-MB-231 cells. We also thank Y. Kurashima, H. Kiyono, M. Kikkawa, M. Aoki, A. Nakajima, E. Tosti, and L.S. Lopez for technical advice. This work was supported in part by Grants-in-Aid for Scientific Research (B) 17H04069 (to HN); Young Scientists (B) 17K15626 (to TN); Grants-in-Aid for Scientific Research (C) 26460397 (to TN); Challenging Exploratory Research 17K19533 (to HN) from Japan Society for the Promotion of Science (JSPS); Scientific Research on Innovative Areas 26110003 (to HN); the Japan Agency for Medical Research and Development (AMED) through AMED-CREST (JP19gm1210002, to HN); Private University Research Branding project (to TM and HN) from the MEXT (Ministry of Education, Culture, Sports, Science and Technology), Japan; and research grants from the Uehara Memorial Foundation (to TN) and the Takeda Science Foundation (to TN).

## AUTHOR CONTRIBUTIONS

T.N. and H.N. designed research; T.N., Y.D., W.T., M.O., D.O., S.Y., M.K., E.N., Y.K., S.A-A., M.H., N.I., and N.T. performed research; M.N., M.O., H.Y., K.S., and T.M. contributed to new reagents/analytical tools; T.N., Y.D., W.T., D.O., N.I., and H.N. analyzed data; T.N., N.I., and H.N. wrote the paper.

## REFERENCES

1. Peterson LW, Artis D. Intestinal epithelial cells: regulators of barrier function and immune homeostasis. Nat Rev Immunol 14, 141–153 (2014).

2. Leoni G, Neumann PA, Sumagin R, Denning TL, Nusrat A. Wound repair: role of immune-epithelial interactions. Mucosal Immunol 8, 959–968 (2015).

3. Kurashima Y, Kiyono H. Mucosal Ecological Network of Epithelium and Immune Cells for Gut Homeostasis and Tissue Healing. Annu Rev Immunol 35, 119–147 (2017).

4. Powell DW, Pinchuk IV, Saada JI, Chen X, Mifflin RC. Mesenchymal cells of the intestinal lamina propria. Annu Rev Physiol 73, 213–237 (2011).

5. Nowarski R, Jackson R, Flavell RA. The Stromal Intervention: Regulation of Immunity and Inflammation at the Epithelial-Mesenchymal Barrier. Cell 168, 362–375 (2017).

6. Isella C, et al. Stromal contribution to the colorectal cancer transcriptome. Nat Genet 47, 312–319 (2015).

7. Calon A, et al. Stromal gene expression defines poor-prognosis subtypes in colorectal cancer. Nat Genet 47, 320–329 (2015).

8. Koliaraki V, Pallangyo CK, Greten FR, Kollias G. Mesenchymal Cells in Colon Cancer. Gastroenterology 152, 964–979 (2017).

9. Smillie CS, et al. Intra- and Inter-cellular Rewiring of the Human Colon during Ulcerative Colitis. Cell 178, 714–730.e722 (2019).

10. Su S, et al. CD10(+)GPR77(+) Cancer-Associated Fibroblasts Promote Cancer Formation and Chemoresistance by Sustaining Cancer Stemness. Cell 172, 841–856 e816 (2018).

11. Putoczki T, Ernst M. More than a sidekick: the IL-6 family cytokine IL-11 links inflammation to cancer. J Leukoc Biol 88, 1109–1117 (2010).

12. Jones SA, Jenkins BJ. Recent insights into targeting the IL-6 cytokine family in inflammatory diseases and cancer. Nat Rev Immunol 18, 773–789 (2018).

13. West NR. Coordination of Immune-Stroma Crosstalk by IL-6 Family Cytokines. Front Immunol 10, 1093 (2019).

14. Gibson DL, et al. Interleukin-11 reduces TLR4-induced colitis in TLR2-deficient mice and restores intestinal STAT3 signaling. Gastroenterology 139, 1277–1288 (2010).

15. Kiessling S, et al. Functional expression of the interleukin-11 receptor alpha-chain and evidence of antiapoptotic effects in human colonic epithelial cells. J Biol Chem 279, 10304–10315 (2004).

16. Yoshizaki A, Nakayama T, Yamazumi K, Yakata Y, Taba M, Sekine I. Expression of interleukin (IL)-11 and IL-11 receptor in human colorectal adenocarcinoma: IL-11 up-regulation of the invasive and proliferative activity of human colorectal carcinoma cells. Int J Oncol 29, 869–876 (2006).

17. Putoczki TL, et al. Interleukin-11 is the dominant IL-6 family cytokine during gastrointestinal tumorigenesis and can be targeted therapeutically. Cancer Cell 24, 257–271 (2013).

18. Molet S, et al. IL-17 is increased in asthmatic airways and induces human bronchial fibroblasts to produce cytokines. J Allergy Clin Immunol 108, 430–438 (2001).

19. Andoh A, et al. Interleukin-22, a member of the IL-10 subfamily, induces inflammatory responses in colonic subepithelial myofibroblasts. Gastroenterology 129, 969–984 (2005).

20. Bamba S, Andoh A, Yasui H, Makino J, Kim S, Fujiyama Y. Regulation of IL-11 expression in intestinal myofibroblasts: role of c-Jun AP-1- and MAPK-dependent pathways. Am J Physiol Gastrointest Liver Physiol 285, G529–538 (2003).

21. Tang W, Yang L, Yang YC, Leng SX, Elias JA. Transforming growth factor-beta stimulates interleukin-11 transcription via complex activating protein-1-dependent pathways. J Biol Chem 273, 5506–5513 (1998).

22. Nishina T, et al. Interleukin-11 links oxidative stress and compensatory proliferation. Sci Signal 5, ra5 (2012).

23. Nishina T, et al. Critical Contribution of Nuclear Factor Erythroid 2-related Factor 2 (NRF2) to Electrophile-induced Interleukin-11 Production. J Biol Chem 292, 205–216 (2017).

24. Calon A, et al. Dependency of colorectal cancer on a TGF-beta-driven program in stromal cells for metastasis initiation. Cancer Cell 22, 571–584 (2012).

25. Zhang X, et al. IL-11 Induces Th17 Cell Responses in Patients with Early Relapsing-Remitting Multiple Sclerosis. J Immunol 194, 5139–5149 (2015).

26. Klein W, et al. A polymorphism in the IL11 gene is associated with ulcerative colitis. Genes Immun 3, 494–496 (2002).

27. Sabzevary-Ghahfarokhi M, et al. The expression analysis of Fra-1 gene and IL-11 protein in Iranian patients with ulcerative colitis. BMC Immunol 19, 17 (2018).

28. Strikoudis A, et al. Modeling of Fibrotic Lung Disease Using 3D Organoids Derived from Human Pluripotent Stem Cells. Cell Rep 27, 3709–3723 e3705 (2019).

29. Schafer S, et al. IL-11 is a crucial determinant of cardiovascular fibrosis. Nature 552, 110–115 (2017).

30. Neufert C, Becker C, Neurath MF. An inducible mouse model of colon carcinogenesis for the analysis of sporadic and inflammation-driven tumor progression. Nat Protoc 2, 1998–2004 (2007).

31. Tanaka T, Kohno H, Suzuki R, Yamada Y, Sugie S, Mori H. A novel inflammation-related mouse colon carcinogenesis model induced by azoxymethane and dextran sodium sulfate. Cancer Sci 94, 965–973 (2003).

32. Moser AR, Pitot HC, Dove WF. A Dominant Mutation That Predisposes to Multiple Intestinal Neoplasia in the Mouse. Science 247, 322–324 (1990).

33. Su LK, et al. Multiple Intestinal Neoplasia Caused by a Mutation in the Murine Homolog of the Apc Gene. Science 256, 668–670 (1992).

34. Deguchi Y, et al. Generation of and characterization of anti-IL-11 antibodies using newly established Il11-deficient mice. Biochem Biophys Res Commun 505, 453–459 (2018).

35. Sakai E, et al. Combined Mutation of Apc, Kras, and Tgfbr2 Effectively Drives Metastasis of Intestinal Cancer. Cancer Res 78, 1334–1346 (2018).

36. Pickert G, et al. STAT3 links IL-22 signaling in intestinal epithelial cells to mucosal wound healing. J Exp Med 206, 1465–1472 (2009).

37. Yoshida GJ, Azuma A, Miura Y, Orimo A. Activated Fibroblast Program Orchestrates Tumor Initiation and Progression; Molecular Mechanisms and the Associated Therapeutic Strategies. Int J Mol Sci 20, (2019).

38. Turley SJ, Cremasco V, Astarita JL. Immunological hallmarks of stromal cells in the tumour microenvironment. Nat Rev Immunol 15, 669–682 (2015).

39. Dasch JR, Pace DR, Waegell W, Inenaga D, Ellingsworth L. Monoclonal antibodies recognizing transforming growth factor-beta. Bioactivity neutralization and transforming growth factor beta 2 affinity purification. J Immunol 142, 1536-1541 (1989).

40. Nandurkar HH, Robb L, Tarlinton D, Barnett L, Kontgen F, Begley CG. Adult mice with targeted mutation of the interleukin-11 receptor (IL11Ra) display normal hematopoiesis. Blood 90, 2148–2159 (1997).

41. Kimura Y, Yanagimachi R. Intracytoplasmic sperm injection in the mouse. Biol Reprod 52, 709–720 (1995).

42. Viennois E, Chen F, Laroui H, Baker MT, Merlin D. Dextran sodium sulfate inhibits the activities of both polymerase and reverse transcriptase: lithium chloride purification, a rapid and efficient technique to purify RNA. BMC Res Notes 6, 360 (2013).

43. Huang DW, Sherman BT, Lempicki RA. Bioinformatics enrichment tools: paths toward the comprehensive functional analysis of large gene lists. Nucleic Acids Research 37, 1–13 (2009).

44. Huang DW, Sherman BT, Lempicki RA. Systematic and integrative analysis of large gene lists using DAVID bioinformatics resources. Nature Protocols 4, 44–57 (2009).

45. Lochner M, et al. Microbiota-induced tertiary lymphoid tissues aggravate inflammatory disease in the absence of RORgamma t and LTi cells. J Exp Med 208, 125–134 (2011).

46. Sato T, et al. Long-term expansion of epithelial organoids from human colon, adenoma, adenocarcinoma, and Barrett’s epithelium. Gastroenterology 141, 1762–1772 (2011).

47. Mihara E, et al. Active and water-soluble form of lipidated Wnt protein is maintained by a serum glycoprotein afamin/alpha-albumin. Elife 5, (2016).

48. Aaltonen LA, Hamilton SR, World Health Organization., International Agency for Research on Cancer. Pathology and genetics of tumours of the digestive system. IARC Press; Oxford University Press (distributor (2000).

49. Sobin LH, Gospodarowicz MK, Wittekind C, International Union against Cancer. TNM classification of malignant tumours, 7th edn. Wiley-Blackwell (2010).

