## Supplementary Materials for "Interleukin-11 is a Marker for Both Cancer- and Inflammation-Associated Fibroblasts that Contribute to Colorectal Cancer Progression"

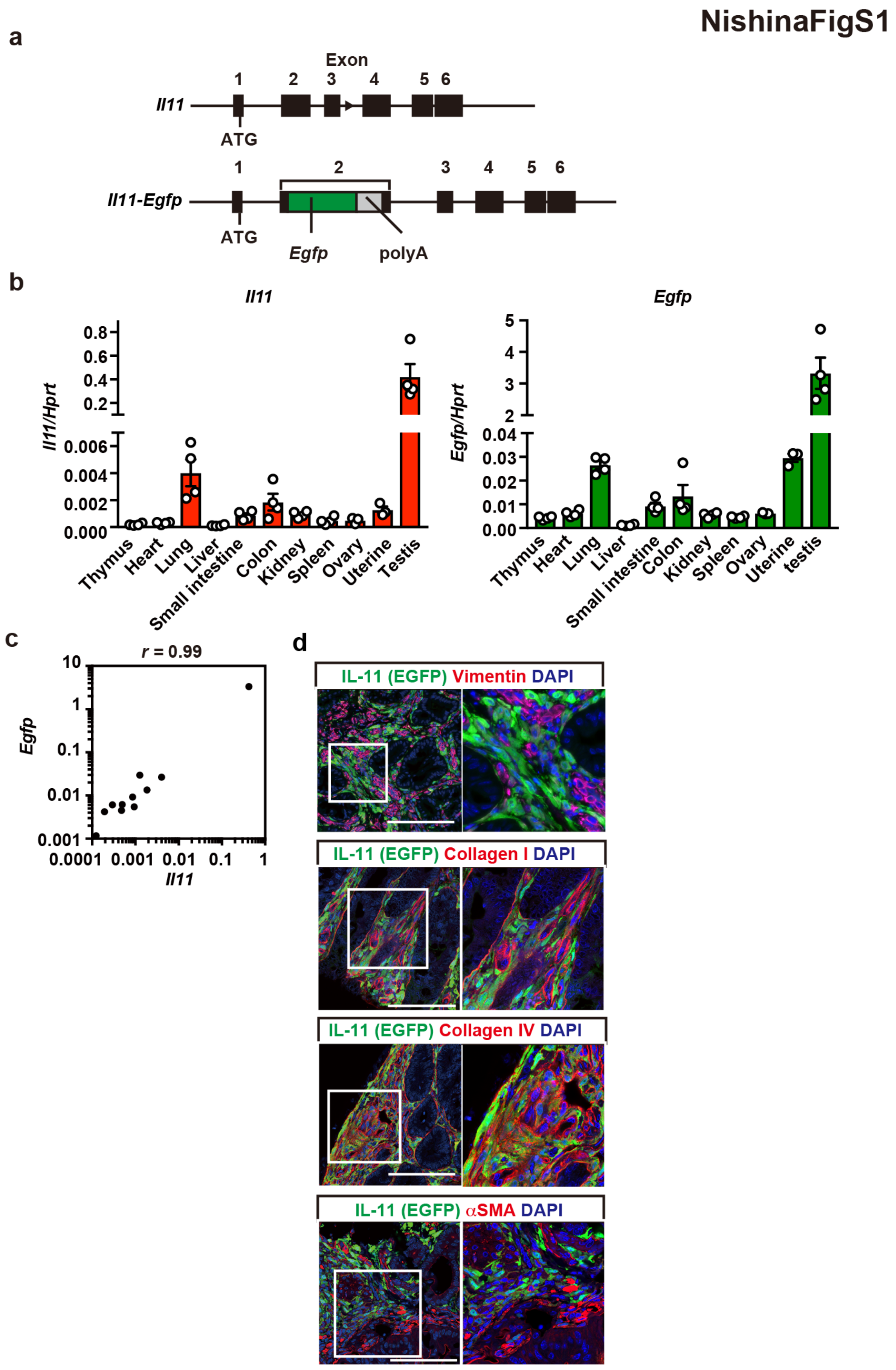

NishinaFigS1
a
Exon
1 2 3 4 5 6
Il11
ATG
1 2 3 4 5 6
Il11-Egfp
ATG
Egfp
polyA
b
Il11
Il11/Hprt
0.8
0.6
0.4
0.2
0.006
0.004
0.002
0.000
Thymus
Heart
Lung
Liver
Small intestine
Colon
Kidney
Spleen
Ovary
Uterine
Testis
Egfp
Egfp/Hprt
5
4
3
2
0.04
0.03
0.02
0.01
0.00
Thymus
Heart
Lung
Liver
Small intestine
Colon
Kidney
Spleen
Ovary
Uterine
testis
c
r = 0.99
Egfp
10
1
0.1
0.01
0.001
0.0001 0.001 0.01 0.1 1
Il11
d
IL-11 (EGFP) Vimentin DAPI
IL-11 (EGFP) Collagen I DAPI
IL-11 (EGFP) Collagen IV DAPI
IL-11 (EGFP) αSMA DAPI

**Supplementary Fig. 1.** Characterization of IL-11$^+$ Cells in CAC Using *Il11-Egfp* Reporter Mice. **a** Diagram of the vector for *Il11-Egfp* reporter mice. The upper panel is the organization of the murine *Il11* gene. An in-frame insertion of *Egfp-polyA* cassette into the second exon of the *Il11* gene resulted in the expression of EGFP with N-terminal three additional amino acids (lower panel). **b, c** *Il11* and *Egfp* expressions in various tissues of *Il11-Egfp* reporter mice. Total RNAs were prepared from the indicated tissues of mice, and *Il11* and *Egfp* expressions were determined by qPCR (**b**). Results are mean ± SE (n = 4 mice). Correlation of *Egfp* and *Il11* expressions (**c**). *Egfp* and *Il11* expression levels were plotted at the X and Y axis, respectively, and the correlation coefficient (*r*) was calculated by the Pearson correlation test. ****p < 0.0001. **d** *Il11-Egfp* reporter mice were treated with AOM/DSS as in Figure 1A, colonic tissue sections were immunostained with the indicated antibodies (red) along with anti-GFP antibody (green). Results are merged images, and the right panels are enlarged images of the boxes (n = 3–4 mice). Scale bar, 100 μm.

NishinaFigS2

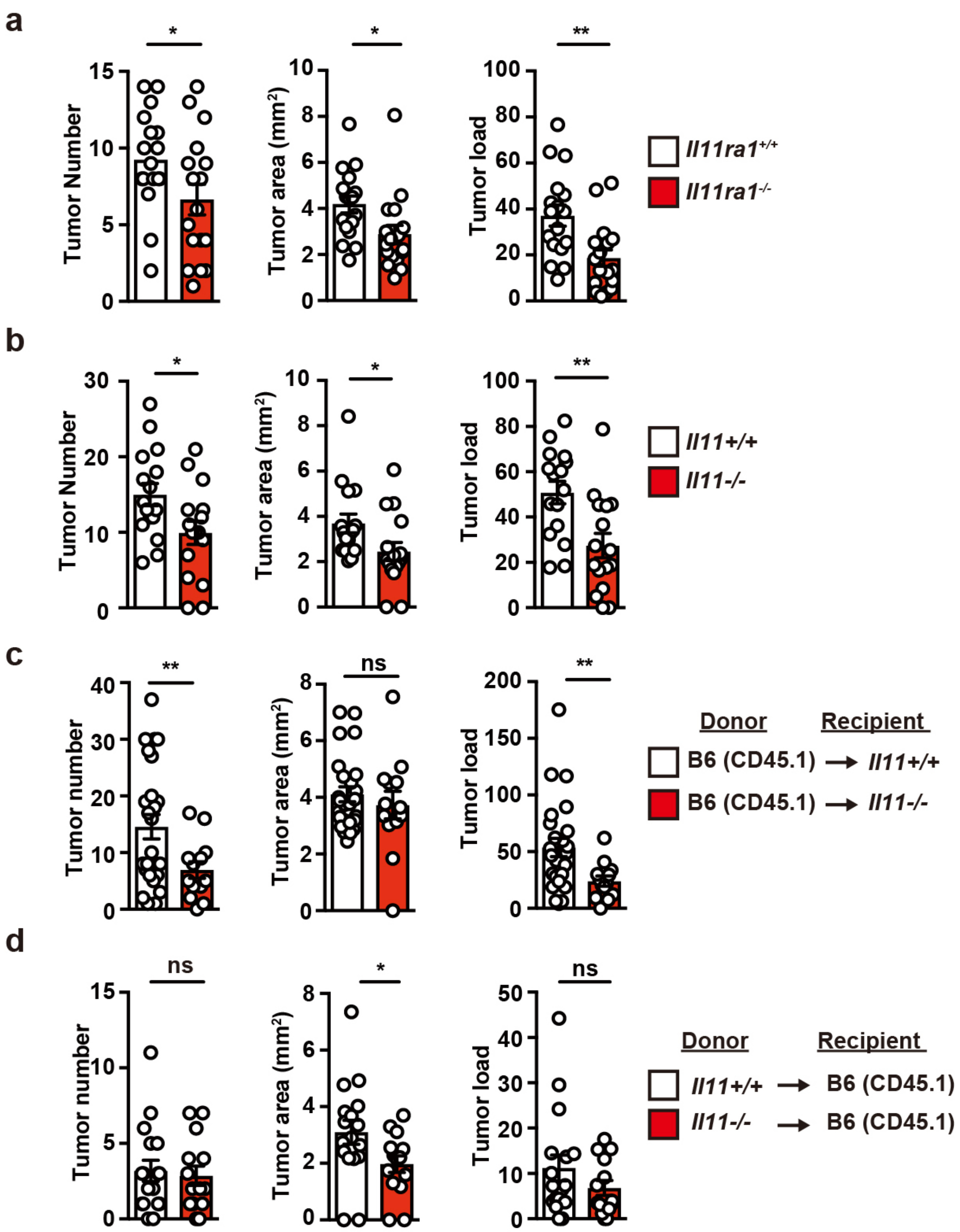


**Supplementary Fig. 2.** Attenuated CAC Development in *Il11ra1*$^{-/-}$ and *Il11*$^{-/-}$ Mice. **a** *Il11ra1*$^{+/+}$ and *Il11ra1*$^{-/-}$ mice were treated as in Figure 1A. On day 84 to 98 after AOM/DSS treatment, mice were sacrificed, and tumor numbers and areas were determined. Tumor load was calculated as in methods, and tumor numbers, average tumor area, and tumor load of an individual mouse are shown. Results are mean ± SE (n = 16 mice). **b** *Il11*$^{+/+}$ and *Il11*$^{-/-}$ mice were treated as in Figure 1A and analyzed as in Figure S2A. Results are mean ± SE (n = 16 mice). **c, d** Deletion of *Il11* in non-hematopoietic cells attenuates development of CAC in mice. BM cells from mice of the indicated genotypes were prepared and transferred into the indicated mice that had been lethally irradiated. At 2 months after BM transfer, mice were treated as in Fig. 1a and analyzed as in Supplementary Fig. 2a. Results are mean ± SE (**d**, n = 13−25 mice; **e**, n = 14−17 mice). Statistical significance was determined by two-tailed unpaired Student's *t*-test (**a−d**). *p < 0.05; **p < 0.01; ***p < 0.001; ns, not significant.

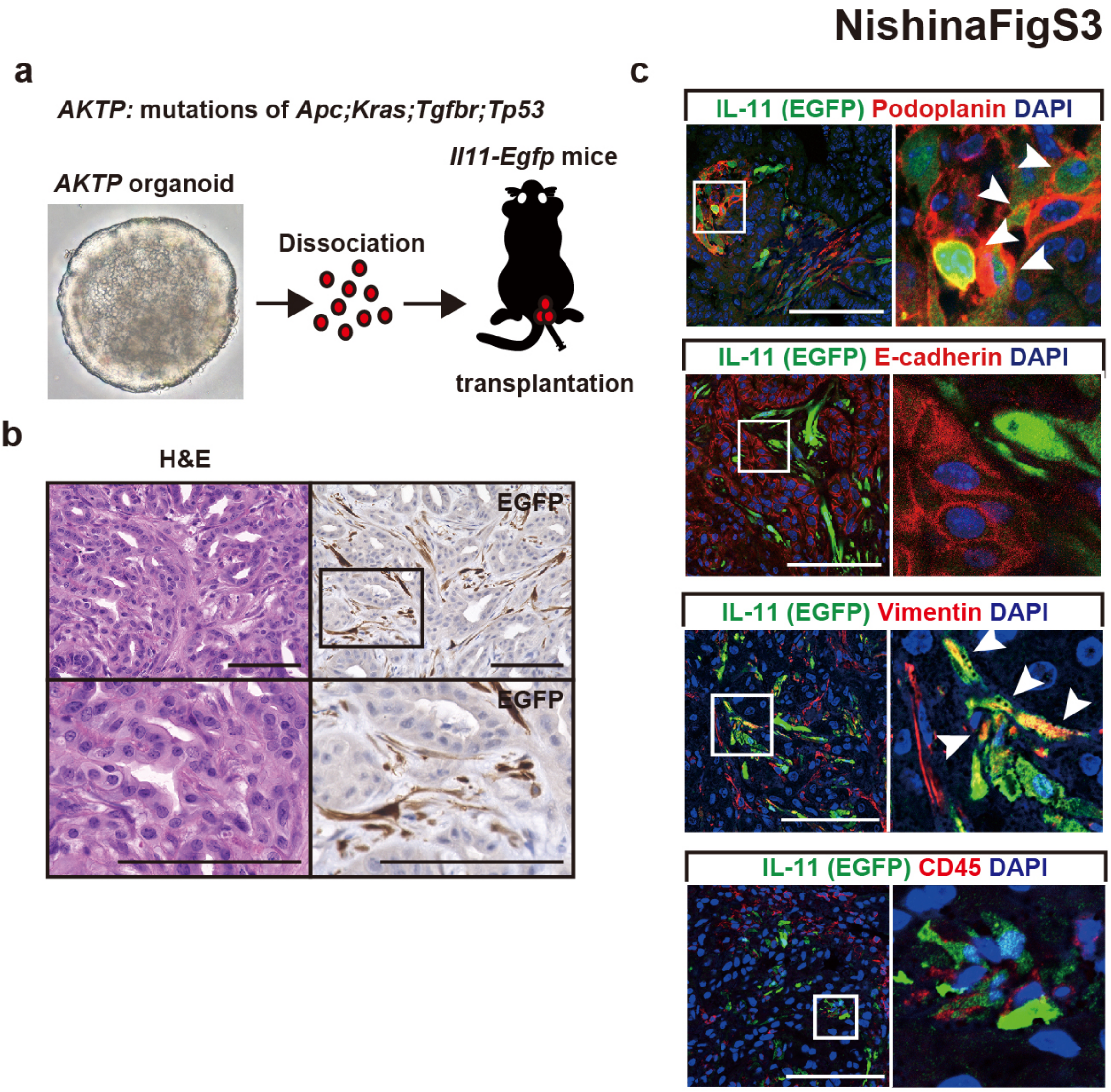


**Supplementary Fig. 3.** Transplantation of Tumor Organoids Causes IL-11$^+$ cell development. **a** Schema of transplantation of the tumor organoids derived from AKTP mice. The tumor organoids from AKTP mice were expanded as a standard procedure. After dissociation, cells were transplanted into the colon of *Il11-Egfp* reporter mice. **b** Tumor organoids develop large tumors, and IL-11$^+$ cells appear in the tumor. Colonic tumor sections were stained with H&E or anti-GFP antibody (B). Scale bar, 100 μm. **c** Tumor sections were stained with anti-E-cadherin, anti-podoplanin, anti-vimentin, or anti-CD45 antibodies (red) along with anti-GFP antibody (green) (n = 4) (**c**). White arrowheads indicate merged cells. Scale bar, 100 μm.

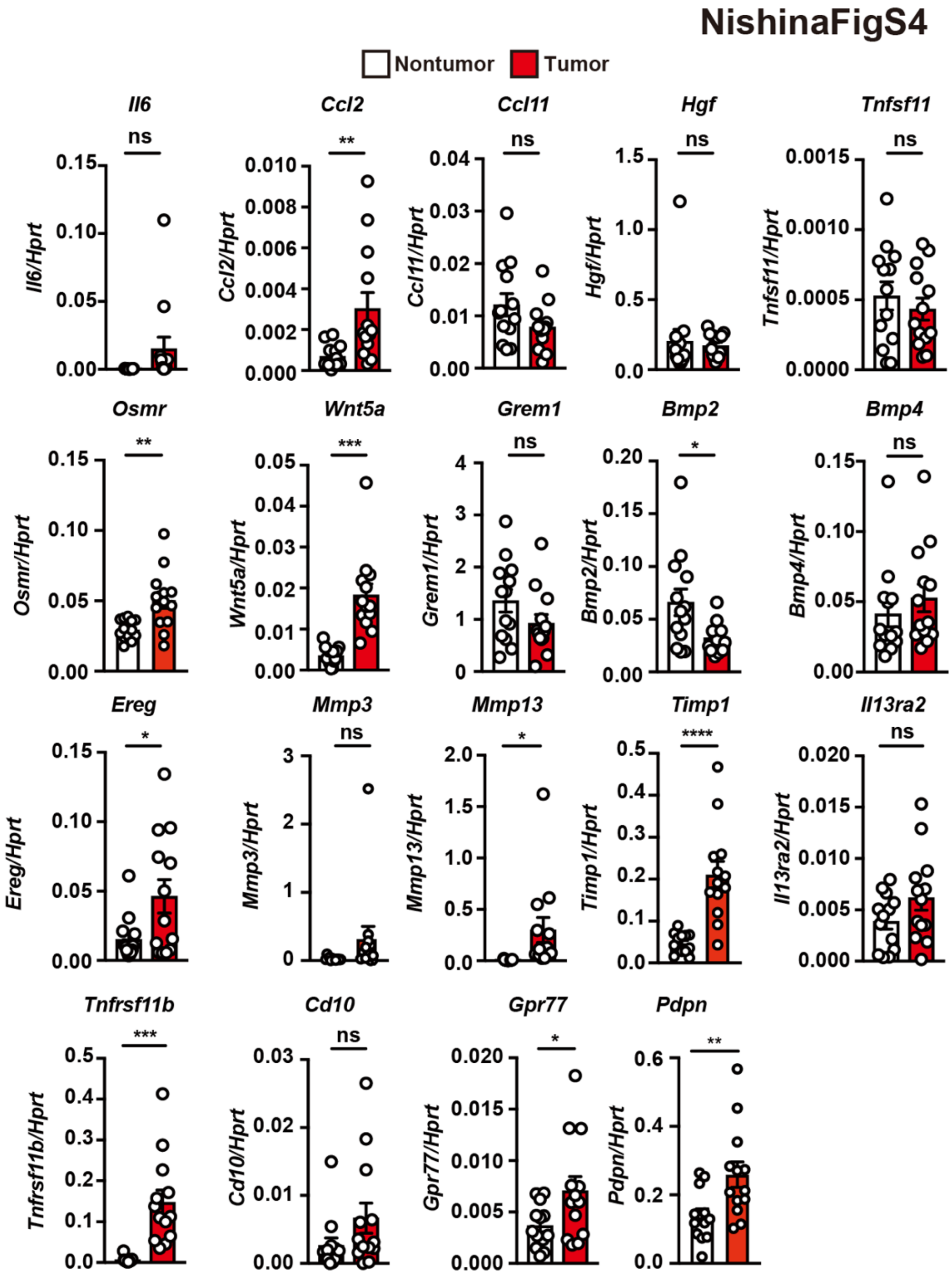


**Supplementary Fig. 4.** Enriched Genes in IL-11$^{+}$ Cells are Elevated in Colon Tumor Tissues of Mice. *Il11-Egfp* reporter mice were treated as in Figure 1A. mRNAs were prepared from tumors and nontumor tissues in the colon of mice on day 98−105 after AOM/DSS treatment, and the expression of the indicated genes was determined by qPCR. Results are mean ± SE (n = 6–10 mice). Statistical significance was determined by two-tailed unpaired Student's *t*-test. *p < 0.05; **p < 0.01; ***p < 0.001; ns, not significant.

**NishinaFigS5**

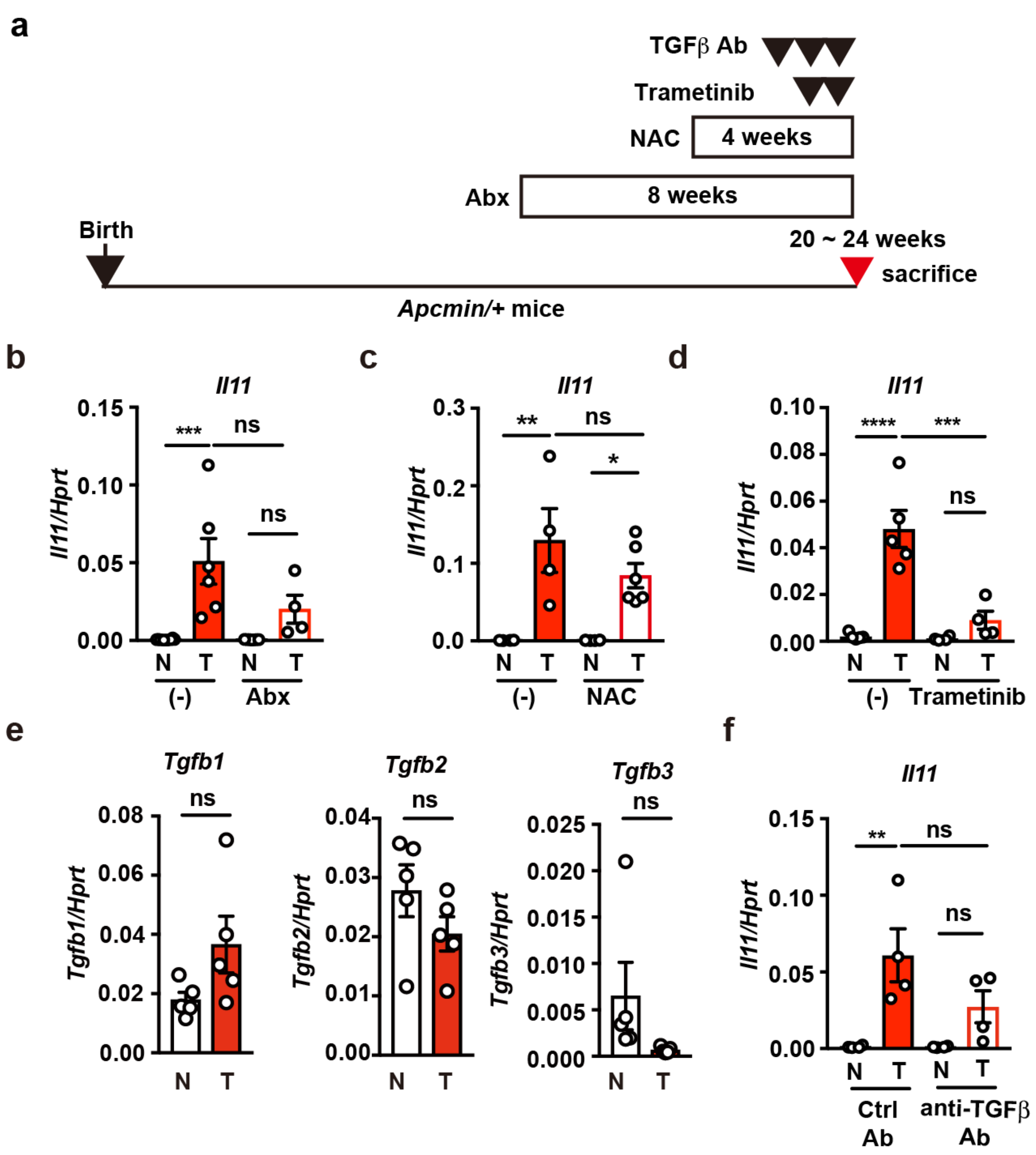


**Supplementary Fig. 5.** The MEK/ERK Pathway is Involved in *Il11* Upregulation in Tumor Tissues of *Apc*$^{min/+}$ Mice. **a** Schema of administration of various inhibitors on *Apc*$^{min/+}$ mice. Mice were treated with Abx (8 weeks), NAC (4 weeks), trametinib (−6 and −30 hours), or anti-TGFβ antibody (days −1, −3, −5) just before sacrifice. **b–i** *Apc*$^{min/+}$ mice were treated with Abx (**b**) (n = 4–8 mice), NAC (**c**) (n = 4–6 mice), or trametinib (**d**) (n = 5 mice) as in A, then sacrificed at 20 to 24 weeks after birth. *Il11* expression in tumors and nontumor tissues in the colon was determined by qPCR. Results are mean ±

SE. **e** *Tgfbs* expression in tumors and nontumor tissues in the colon of *Apc*$^{min/+}$ mice was determined by qPCR. Results are mean ± SE (n = 5 mice). **f** *Apc*$^{min/+}$ mice were intraperitoneally injected with control mouse IgGs or anti-TGFβ antibody as in A. *Il11* expression in tumors and nontumor tissues was determined by qPCR. Results are mean ± SE (n = 4 mice). Statistical significance was determined by two-way ANOVA with Bonferroni's test (**b, c, d, f**), or two-tailed unpaired Student's *t*-test (**e**). *p < 0.05; **p < 0.01; ***p < 0.001; ****p < 0.0001; ns, not significant.

NishinaFigS6

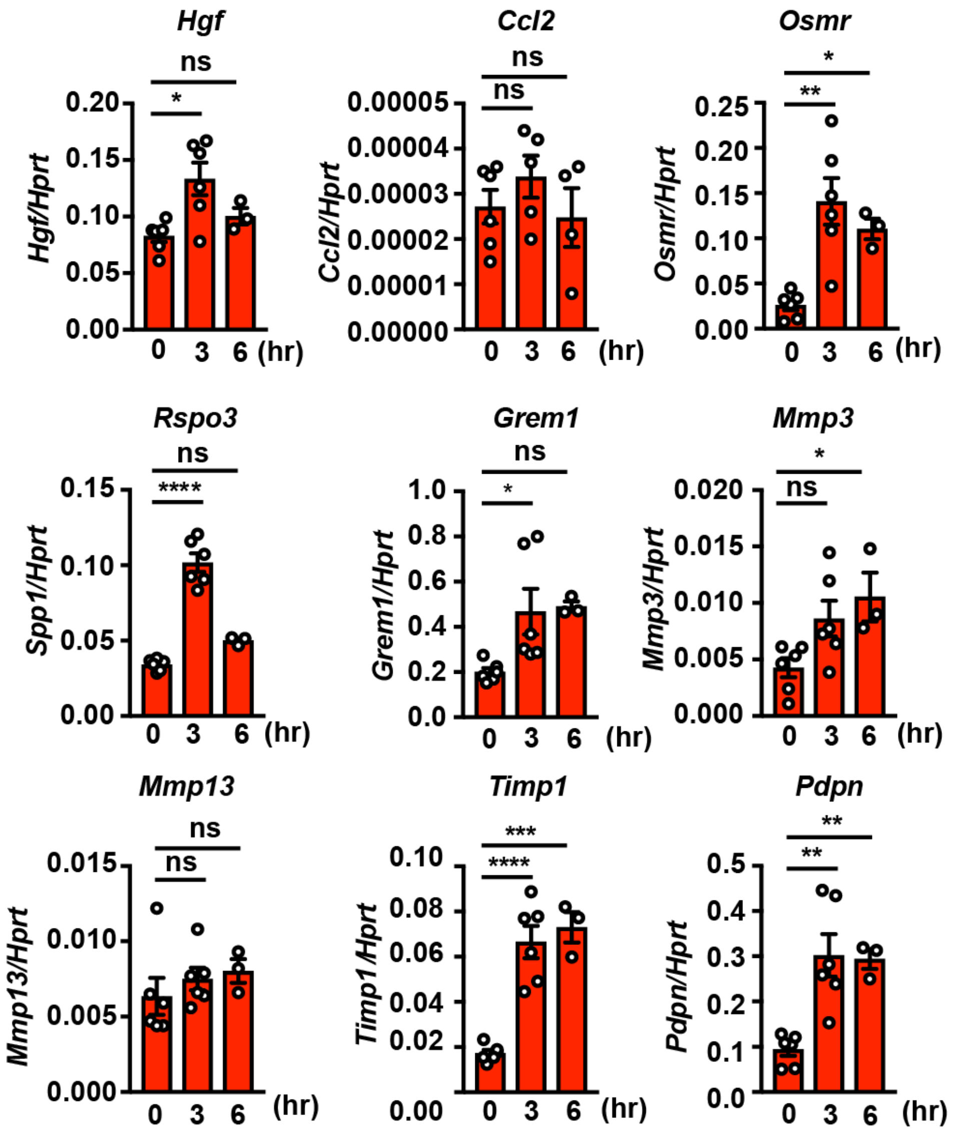


**Supplementary Fig. 6.** Administration of IL-11R Agonist Induces the Expression of Genes Enriched in IL-11$^+$ cells. **a, b** Wild-type mice were treated with IL-11R agonist as in Fig. 6c, the expression of the indicated genes at the indicated times after injection was determined by qPCR. Results are mean ± SE (n = 3−6 mice). The expression of genes enriched in IL-11$^+$ cells are shown. Statistical significance was determined by one-way ANOVA with Tukey's post-hoc test. *p <0 .05; **p < 0.01; ***p < 0.001; ****p < 0.0001; ns, not significant,

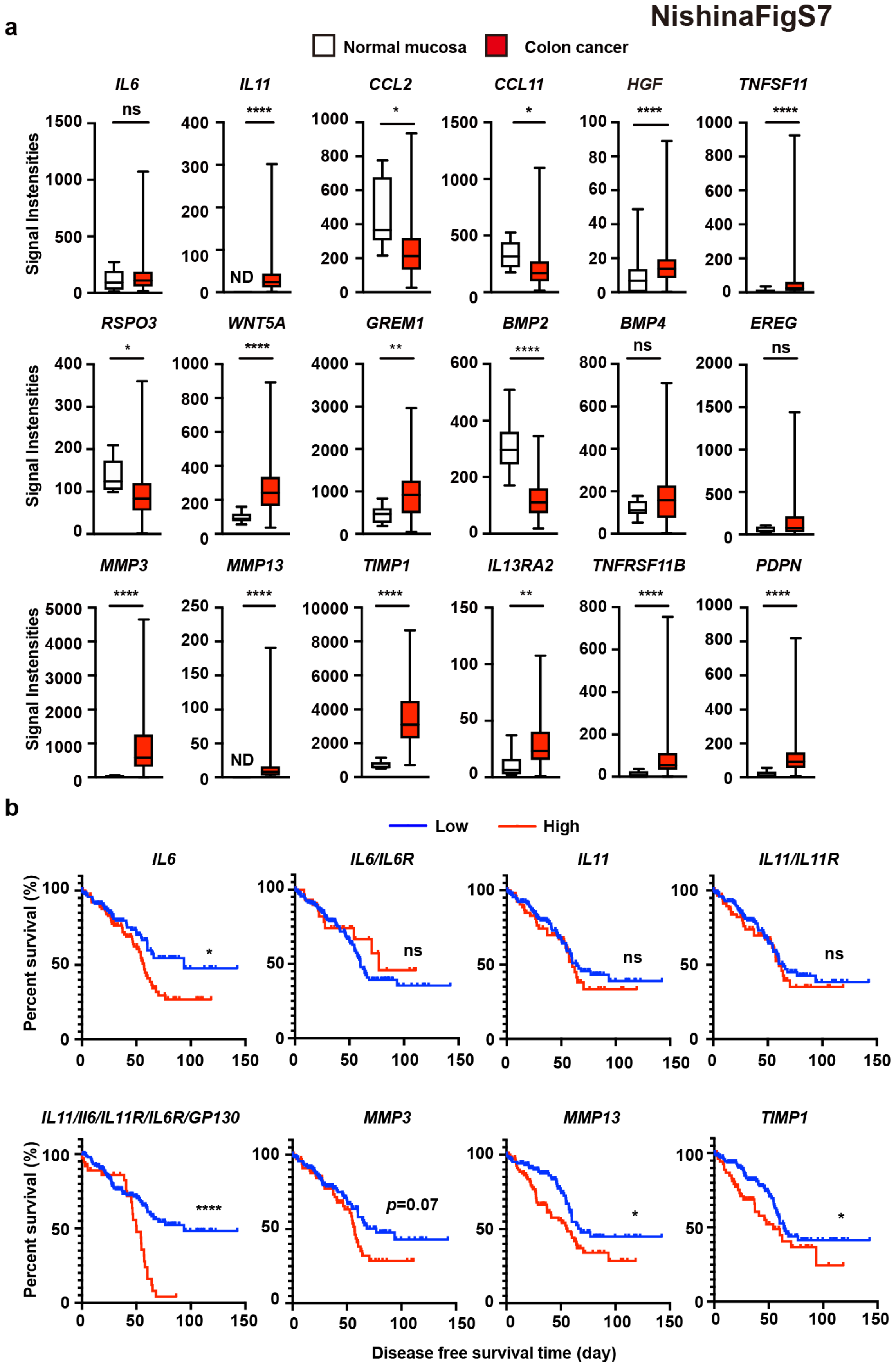
NishinaFigS7
a
Normal mucosa
Colon cancer
Signal Instensities
IL6
IL11
CCL2
CCL11
HGF
TNFSF11
RSPO3
WNT5A
GREM1
BMP2
BMP4
EREG
MMP3
MMP13
TIMP1
IL13RA2
TNFRSF11B
PDPN
ns
*
**
****
ND
b
Low
High
Percent survival (%)
IL6
IL6/IL6R
IL11
IL11/IL11R
IL11/Il6/IL11R/IL6R/GP130
MMP3
MMP13
TIMP1
p=0.07
Disease free survival time (day)

**Supplementary Fig. 7.** Genes Enriched in IL-11$^+$ IAFs are Elevated in Human Colorectal Cancer. **a** Genes enriched in IL-11$^+$ IAFs are elevated in human colon cancer tissues. From the data set (GSE33113), we retrieved the expressions of the genes enriched in IL-11$^+$ IAFs. The signaling intensities of each gene in normal mucosa (n = 6) and colon cancer tissues (n = 90) are shown. Results are mean ± SE. Statistical significance was determined by Mann-Whitney U test. *p < 0.05; **p < 0.01; ****p < 0.0001; ns, not significant. **b** High expression of *IL6*, the combination of *IL6/IL6R/IL11/IL11R/GP130*, *MMP13*, and *TIMP1* were correlated with reduced disease-free survival in colon cancer patients. GSE17536 and GSE17537 were obtained from the Gene Expression Omnibus. For the genes enriched in IL-11$^+$ IAFs, we classified the expression levels in these data sets into high and low using hierarchical clustering. Then each gene enriched in IL-11$^+$ IAFs was correlated with survival using the Kaplan-Meier method (n = 232). Statistical significance was determined by Mantel-Cox log-rank test. *p < 0.05; ****p < 0.0001; ns, not significant.

**Table S1. List of primers used in this study.**

| Target Gene | Forward Primer | Reverse Primer |
|---|---|---|
| *16S rRNA* | GGT GAA TAC GTT CCC GG | TAC GGC TAC CTT GTT ACG ACT T |
| *Bmp2* | GGG ACC CGC TGT CTT CTA GT | TCA ACT CAA ATT CGC TGA GGA C |
| *Bmp4* | TTC CTG GTA ACC GAA TGC TGA | CCT GAA TCT CGG CGA CTT TTT |
| *Ccl2* | CCC AAT GAG TAG GCT GGA GA | AAA ATG GAT CCA CAC CTT GC |
| *Ccl11* | GAA TCA CCA ACA ACA GAT GCA C | ATC CTG GAC CCA CTT CTT CTT |
| *Cd10* | CTC TCT GTG CTT GTC TTG CTC | GAC GTT GCG TTT CAA CCA GC |
| *Egfp* | AGC AAA GAC CCC AAC GAG AA | GGC GGC GGT CAC GAA |
| *Ereg* | TGC CTC TTG GGT CTT GAC G | ACT TTG TAA TCT GCA CTT GAG CC |
| *Gpr77* | CTG GGC CTC TTG CTG ACT GTG C | GCC CCA GGA AGC CAA AGA GGA |
| *Grem1* | TCA AAG CGG GCA CAT TCA G | AGT AGG AAT CGG GTG GTT TGG |
| *Hgf* | ATG TGG GGG ACC AAA CTT CTG | GGA TGG CGA CAT GAA GCA G |
| *Hmox1* | GTC AAG CAC AGG GTG ACA GA | ATC ACC TGC AGC TCC TCA AA |
| *Hprt* | AAC AAA GTC TGG CCT GTA TCC AA | GCA GTA CAG CCC CAA AAT GG |
| *Il11* | CTG CAC AGA TGA GAG ACA AAT TCC | GAA GCT GCA AAG ATC CCA ATG |
| *Il11ra1* | AAC AGA TGC TGT GGC TGG G | CAG GGG ACC AGT GCT AGG AG |
| *Il13ra2* | ACC GAA ATG TTG ATA GCG ACA G | ACA ATG CTC TGA CAA ATG CGT A |

| | | |
|---|---|---|
| *Il22ra1* | ATG AAG ACA CTA CTG ACC ATC CT | CAG CCA CTT TCT CTC TCC GT |
| *Il6* | GTA TGA ACA ACG ATG ATG CAC TTG | ATG GTA CTC CAG AAG ACC AGA GGA |
| *Il6st* | CCG TGT GGT TAC ATC TAC CCT | CGT GGT TCT GTT GAT GAC AGT G |
| *Mmp3* | ACA TGG AGA CTT TGT CCC TTT TG | TTG GCT GAG TGG TAG AGT CCC |
| *Mmp13* | CTT CTT CTT GTT GAG CTG GAC TC | CTG TGG AGG TCA CTG TAG ACT |
| *Osmr* | TAT TTC TTG GGA GCC CGT AT | TCT GAA GTT GTA ACG GAC GC |
| *Pdpn* | ACC GTG CCA GTG TTG TTC TG | AGC ACC TGT GGT TGT TAT TTT GT |
| *Rspo3* | CCA ACC AGC GAG ACA AGA AC | GAG GAG GAG CTT GTT TCC TTT C |
| *Tgfb1* | TTG CTT CAG CTC CAC AGA GA | TGG TTG TAG AGG GCA AGG AC |
| *Tgfb2* | CTT CGA CGT GAC AGA CGC T | GCA GGG GCA GTG TAA ACT TAT T |
| *Tgfb3* | CAG GCC AGG GCA GTC AGA G | ATT TCC AGC CTA GAT CCT GCC |
| *Timp1* | GCA ACT CGG ACC TGG TCA TAA | CGG CCC GTG ATG AGA AAC T |
| *Tnfrsf11b* | ACC CAG AAA CTG GTC ATC AGC | CTG CAA TAC ACA CAC TCA TCA CT |
| *Tnfsf11* | CAG CAT CGC TCT GTT CCT GTA | CTG CGT TTT CAT GGA GTC TCA |
| *Wnt5a* | CAA CTG GCA GGA CTT TCT CAA | CAT CTC CGA TGC CGG AAC T |
